# Evidence for a rod-to-cone lactate shuttle in the mammalian retina

**DOI:** 10.64898/2026.08.21.744797

**Authors:** Lan Wang, Wadood Haq, Lucia Peiroten, Alexander Hirsch, Catherine Hottin, Laimdota Zizmare, Yiyi Chen, Victor Calbiague, Paul A. Roberts, Oliver Schmachtenberg, Christoph Trautwein, François Paquet-Durand

**Affiliations:** Institute for Ophthalmic Research, University of Tübingen, 72076, Germany; Werner Siemens Imaging Center, Department of Preclinical Imaging and Radiopharmacy, University of Tübingen, 72076, Germany; Core Facility Metabolomics, Faculty of Medicine, University of Tübingen, 72076, Germany; Instituto de Biología y CINV, Facultad de Ciencias, Universidad de Valparaíso, Chile; Institut de la Vision, CNRS, INSERM, Sorbonne University, 75012 Paris, France; Department of Optometry and Visual Science, City St. George’s, University of London, London, United Kingdom; Yunnan Eye Institute & Key Laboratory of Yunnan Province, Yunnan Eye Disease Clinical Medical Center, Affiliated Hospital of Yunnan University, Yunnan University, 176 Qingnian Road, 650021 Kunming, China

**Keywords:** Energy metabolism, retinal degeneration, neurodegeneration, ANLS, *rd1* mouse

## Abstract

In his seminal 1920s studies, Otto Warburg found the retina to generate large amounts of lactate. However, it was unclear what retinal cells produced lactate and whether it was a metabolic waste product or used further. Here, we show that lactate produced by rod photoreceptors fuels the energy-intensive function and viability of cone photoreceptors.

In an initial expression analysis, we found monocarboxylate transporter-1 (MCT1), lactate-producing lactate-dehydrogenase-A (LDHA), and pyruvate carboxykinase-1 (PCK1) localized to rod photoreceptors, while high-affinity MCT2, pyruvate-producing LDHB, and PCK2 were expressed in cones. We then exposed retina to defined media containing either glucose or lactate as caloric component, and applied specific MCT inhibitors. In glucose-containing medium, ^1^H-NMR metabolomics showed rod MCT1 inhibition to increase retinal lactate, suggesting rods as a major source of lactate. In lactate-only medium, functional recordings using micro-electroretinography showed decreased rod function, while cone function was maintained. In glucose-containing medium, blocking rod MCT1 abolished cone function. Long-term treatment with MCT inhibitors selectively decreased photoreceptor viability. Conversely, supplementing the defined medium with lactate preserved cone viability in the *rd1* mouse model for *Retinitis Pigmentosa*.

Together, our data suggest that lactate shuttling from rods is crucial for cone function and viability. This may explain cone degeneration seen in various retinal diseases and provides an entirely new avenue for metabolism-based treatment development. The discovery of a lactate-shuttle between two functionally similar, yet distinct types of neurons may have far-reaching implications for our understanding of the central nervous system in general.

**Highlights:**

- Rod *function* is driven mainly by glycolysis, rod *viability* requires oxidative phosphorylation
- Rod photoreceptors export lactate, cone photoreceptors import lactate
- Cone *function* and *viability* depends on rod glycolysis and cone oxidative phosphorylation
- Inhibition of lactate transport causes photoreceptor degeneration
- Lactate supplementation prevents secondary cone degeneration in *rd1* mouse retina

## Introduction

The retina is the part of the central nervous system (CNS) with the highest per weight energy consumption. The most abundant retinal cell type are the photoreceptors – rods and cones – characterized by an extraordinarily high energy demand, notably to fuel phototransduction^1^. Surprisingly, the energy-hungry retina appears to use primarily energy-inefficient glycolysis for ATP-production, associated with the release of large amounts of lactate^2^. This phenomenon was discovered already in the 1920s by Otto Warburg^3^, yet, to this day it is unclear why in the retina high ATP demand is paired with inefficient production. However, this paradox might be resolved if lactate generated was used by other retinal cell types.

In general, mammalian cells employ glycolysis to split glucose into two pyruvate molecules and generate two moles of ATP. Pyruvate may enter the mitochondrial Krebs-cycle and undergo oxidative phosphorylation (OXPHOS), to produce further 32 moles of ATP per glucose molecule^4^. Thus, ATP generation via OXPHOS is far more efficient per carbon input than glycolysis. To maintain glycolytic flux when pyruvate is not consumed by OXPHOS, pyruvate is reduced to lactate, traditionally seen as a metabolic waste product.

Nevertheless, lactate can be used as fuel for OXPHOS and lactate shuttling can transfer oxidative substrates between different cell types^5^. This notion gave rise to the astrocyte-neuron-lactate-shuttle (ANLS) hypothesis, which proposed that lactate generated through glycolysis in glia cells was transferred to neurons to be used as fuel for Krebs-cycle and OXPHOS^6^. However, CNS lactate shuttling remains controversial^7,8^ and whether and where lactate shuttling occurs in the retina is unclear.

Photoreceptors have been suggested to be the main consumers of glucose and principal producers of lactate, which was suggested to in part fuel retinal pigment epithelial (RPE) cells or Müller glia cells^9^. Recent research indicated significant differences in the metabolism of rod and cone photoreceptors^10,11^, which may relate to the fact that cone photoreceptors expend twice as much energy as rods, do not saturate in bright light, and harbor more and larger synapses with second-order retinal neurons^1^. The fundamental differences between rod and cone energy demand and dynamics likely necessitate distinct metabolic strategies.

Here, we show that the function of cone photoreceptors crucially depends on lactate shuttling from rods. This result is in line with differential expression of key lactate metabolism enzymes in rods and cones, and their metabolic responses to monocarboxylate transporter (MCT) inhibitors. Importantly, rod-to-cone lactate shuttling may explain the secondary loss of cone function seen in retinal diseases, such as *Retinitis Pigmentosa* (RP) and age-related macular degeneration (AMD). This may provide an entirely new avenue for metabolism-based treatment development. Moreover, lactate shuttling between two different types of neurons resolves a long-standing mystery in retinal metabolism and may have significant ramifications for our understanding of CNS functioning as a whole.

## Results

### Retinal expression of transporters and key enzymes related to lactate metabolism

We have previously found the glucose transporters -1 and -3 to be mainly expressed in RPE and rod photoreceptors, respectively^11^. To understand the resulting flow of lactate (lac) within the retina, we first investigated the cell-specific localization of relevant enzymes and transporters.

MCTs are bidirectional lac transporters and MCT1, with a k_m_ value of ≈5 mM, can be expected to be both importer and exporter, depending on lac gradients. In contrast, the low k_m_ value of 0.5 mM for MCT2 means that it is more likely to be a lac importer. MCT1 immunofluorescence revealed strong expression in the inner segments and synapses of rod photoreceptors (Figure 1a). In contrast, in the outer nuclear layer (ONL) MCT2 was specifically localized to cone photoreceptors, as evidenced by co-localization with the cone-specific marker peanut agglutinin (PNA). MCT2 was also abundantly expressed in inner nuclear layer (INL) bipolar cells (Figure 1b). Further immunofluorescence investigations for MCT3 and MCT4 revealed their expression on the basal side of the RPE and in Müller glial cells, respectively (Supplemental Figure S1).

**Figure 1.**
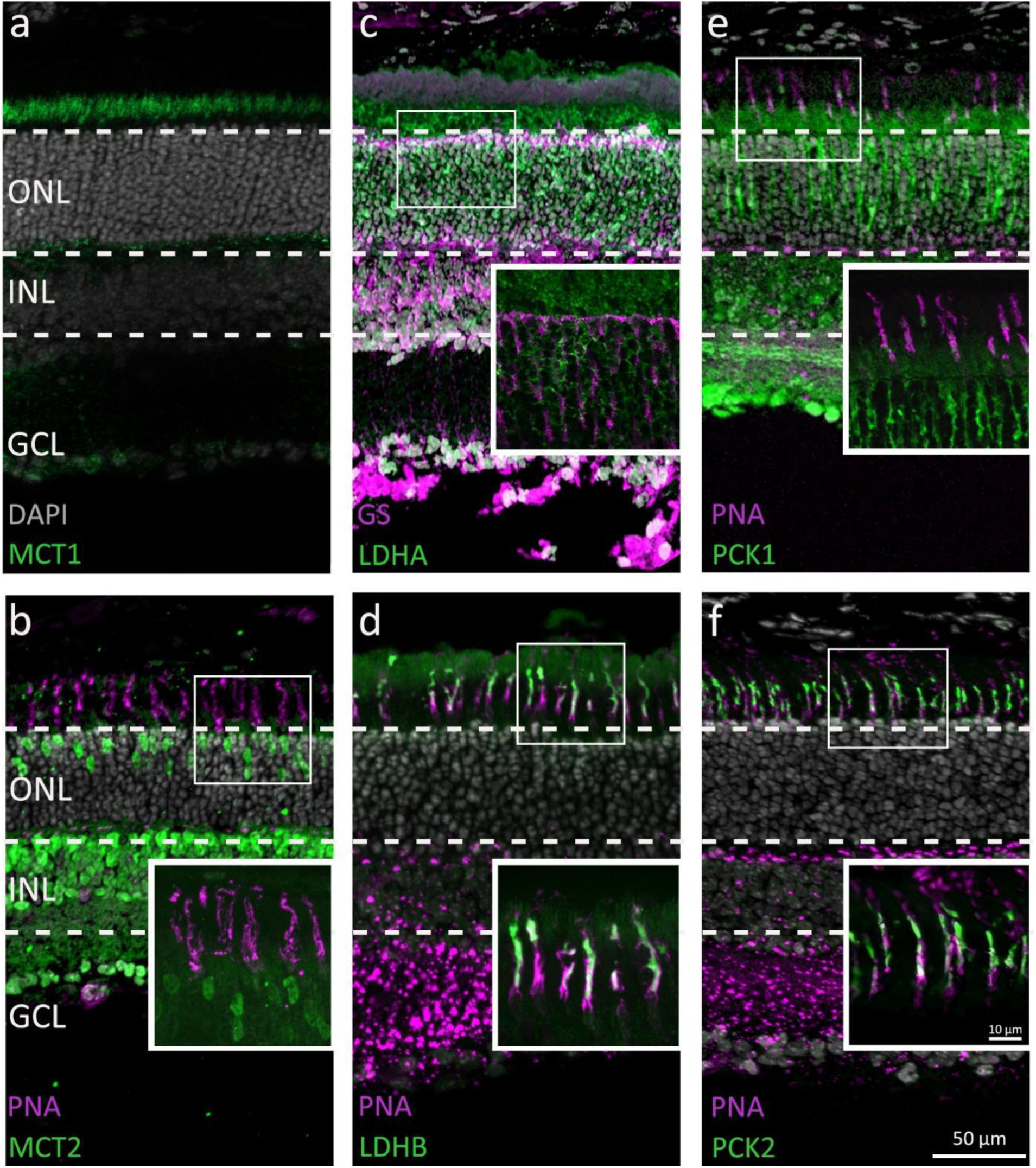
Expression of key enzymes and transporters involved in lactate metabolism. Immunodetection of key metabolic enzymes (green) at post-natal day (P) 30 in wild-type retina. DAPI (grey) was used as a nuclear counterstain. Co-localization with Müller cell marker glutamine synthetase (GS; magenta) and cone photoreceptor marker peanut agglutinin (PNA; magenta) was used to confirm cell-type specific expression. **a)** Monocarboxylate transporter-1 (MCT1) immunolabeling in rod inner segments and synapses. **b)** MCT2 label in cone somata and bipolar cells. **c)** Lactate dehydrogenase A (LDHA) expression in rod photoceptor cell bodies and inner segments. **d)** Immunofluorescence for pyruvate-generating LDHB antibody labels predominantly cone inner segments. **e)** Phosphoenolpyruvate carboxykinase -1 (PCK1) labels somata of rod photoreceptors. **f)** PCK2 labels cone inner segments. ONL=outer nuclear layer, INL=inner nuclear layer, GCL=ganglion cell layer; scale bar=50 µm.

Next, we examined retinal expression of lactate dehydrogenase (LDH), a tetrameric enzyme which catalyzes the conversion of pyruvate (pyr) to lac or *vice versa*^12^. While LDHA preferentially reduces pyr to lac, LDHB catalyzes the oxidation of lac to pyr^13^. LDHA was mainly expressed in the soma and inner segments of rods, whereas strong LDHB expression was found in cone inner segments (Figure 1c, d). Thus, rods possess the enzymatic machinery to produce lac, whereas cones are likely to consume lac as an energy source.

The cytoplasmic phosphoenolpyruvate carboxykinase-1 (PCK1) and the mitochondrial PCK2 convert oxaloacetate (OAA) to phosphoenolpyruvate (PEP) and CO_2_. PCK1 was found to be strongly expressed in rod photoreceptor cytoplasm, while PCK2 was detected in cone inner segments, as shown by PNA co-staining (Figure 1e, f). The dephosphorylation of PEP to pyr is catalyzed by pyruvate kinase M2 (PKM2), an enzyme expressed in photoreceptor inner segments^11^ (*cf*. Supplemental Figure S1).

Taken together, these results show that rods and cones differ markedly in their expression of key enzymes involved in lac and pyr metabolism, with notably the co-expression of MCT1 / LDHA in rods and MCT2 / LDHB in cones, indicating that cones could consume lac released by rods. Moreover, cones within their mitochondria would have the ability to flexibly regulate acetyl-CoA pools for Krebs-cycle activity, using the conversion of OAA to pyr via PCK2 / PKM2.

### Inhibition of lactate shuttling differentially affects photoreceptor survival

To functionally confirm the MCT expression patterns seen above and to delineate the roles of lac transport for photoreceptor energy metabolism and survival, we applied selective MCT inhibitors to organotypic retinal explants cultured in defined, serum-free R16 medium, containing 19.2 mM glc. The effects of different treatments were characterized by quantifying ONL cell death (Figure 2a), as well as the number of surviving cones (Figure 2b). For the MCT inhibitors used – AZD3965 (AZD), a potent, selective MCT1 inhibitor^14^ and AR-C155858 (ARC), a widely used dual inhibitor of MCT1 and MCT2^15^ – dose-response data was collected (Supplemental Figure S2).

**Figure 2.**
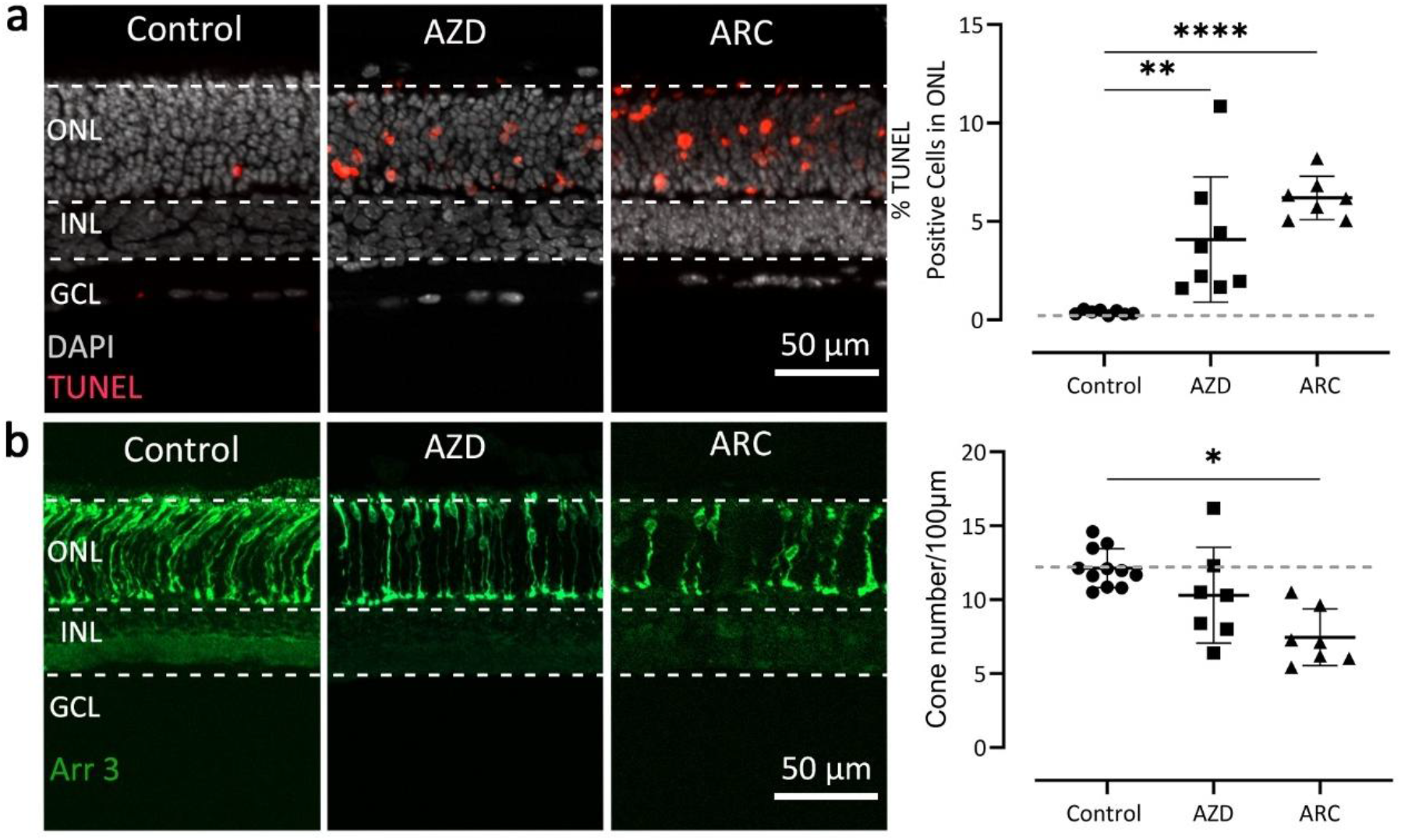
MCT inhibition causes photoreceptor degeneration. **a)** TUNEL assay labelling dying cells (red) in wild-type retinal explant cultures. DAPI (grey) was used as nuclear counterstain. Untreated (Control) retina was compared to retina treated with 5 µM AZD3965 (AZD) or 2 µM AR-C155858 (ARC). The scatter plot shows percentages of TUNEL positive cells in the outer nuclear layer (ONL). **b)** Cone-arrestin labelling (green) in *wt* retinal explant cultures. The scatter plot shows numbers of arrestin-positive cones in the ONL. Statistical testing: One-way ANOVA and Dunnett’s multiple comparisons test. Grey dashed lines designate control conditions, data points above or below this threshold indicate destructive effects; n=6-10 independent retinal explants; error bars represent SD; significance levels: *=p<0.05; **=p<0.01; ****=p<0.0001. INL=inner nuclear layer, GCL=ganglion cell layer; scale bar=50 µm.

In wild-type control retina, the cell death rate, as assessed by the TUNEL assay, was low (0.38%±0.04, n=8) and cone counts were high at 12.14 (±0.40, n=11) cells per 100 µm of retinal circumference. Treatment of wild-type retina with AZD led to a significant increase in photoreceptor cell death (4.10%±1.12, n=8, p<0.01); however, cone cell numbers remained largely unaffected (11.01±0.98, n=7, p=0.99). Treatment with SR-13800, another highly specific MCT1 inhibitor, confirmed these results (Supplemental Figure S3). Since there are currently no MCT2 selective inhibitors available, we employed ARC to probe for MCT2 function. Following ARC treatment, we observed a further increase in ONL cell death (6.20%±0.41, n=7, p<0.0001), concomitant with a significant reduction of cone viability (7.46±0.72, n=7, p<0.05).

Together these experiments showed that blocking lac transport causes photoreceptor degeneration, in line with earlier genetic experiments^16^, but the vulnerabilities of rod and cone photoreceptors differed markedly. MCT1 inhibition primarily affected rods, whereas combined inhibition of MCT1 and MCT2 also caused strong cone loss. Since MCT2 was expressed only on cone photoreceptors, this suggested that cone survival depended on lac import.

### Blocking lactate transport alters retinal metabolic signatures

To evaluate the metabolic alterations of MCT inhibition, we employed organotypic retinal explant cultures and high-field (600 MHz) ^1^H-NMR spectroscopy-based metabolomics. This allowed to identify distinct metabolites in the retina and in the supernatant defined culture medium. In the retina, both AZD- and ARC-treatment produced marked changes in metabolic patterns (Figure 3a, b). Among the 18 significantly altered metabolites was lac, which was strongly elevated after MCT inhibition, supporting the idea that MCT1 was primarily a lac exporter. Pattern hunter analysis showed O-acetylcarnitine, GDP, and ADP as the metabolites most correlated with lac (Figure 3c). Since carnitine can take up acetyl groups from acetyl-CoA and serve as an “acetyl-buffer”, high levels of acetylcarnitine could indicate insufficient Krebs-cycle activity.

**Figure 3.**
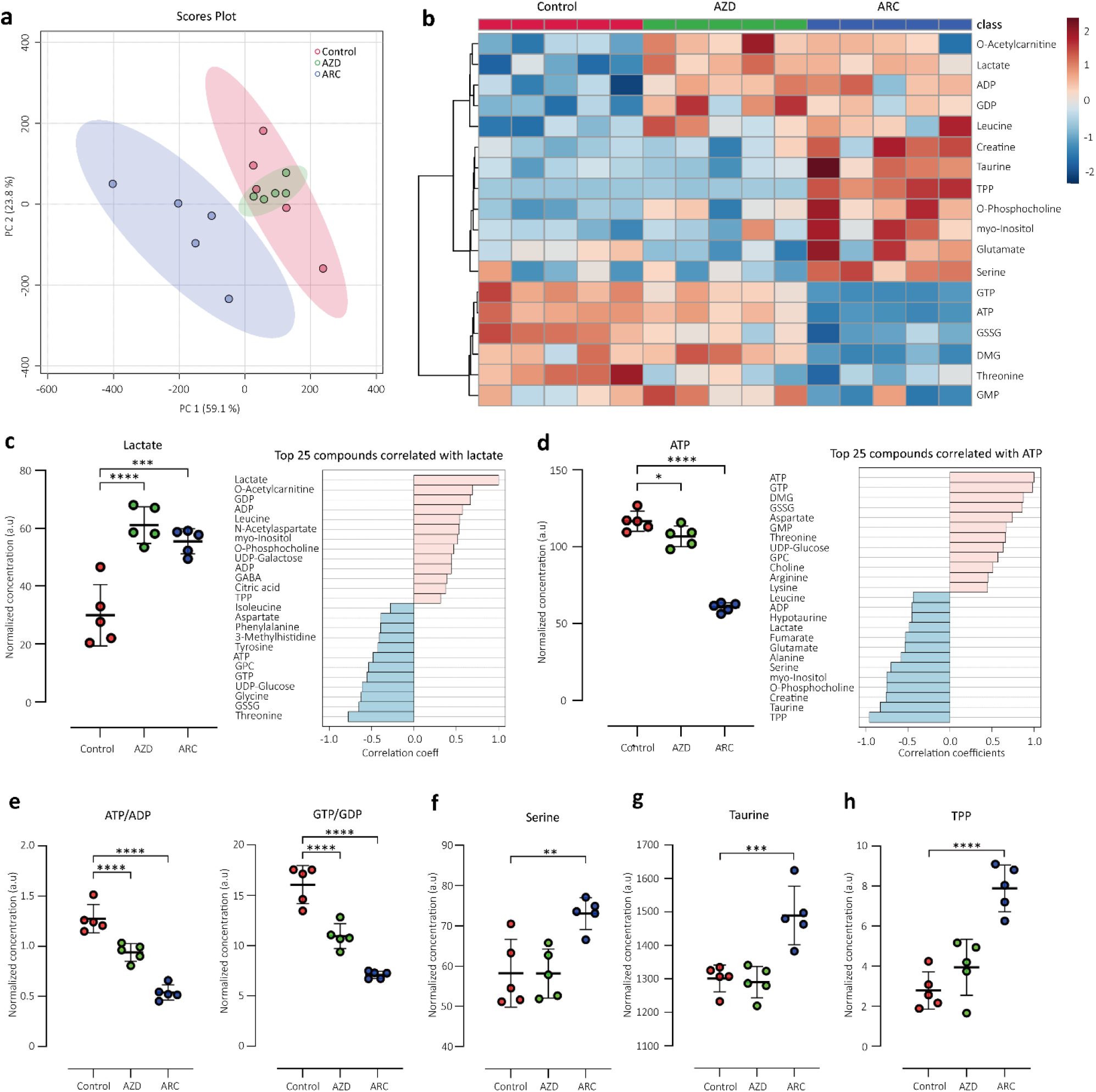
MCT inhibition causes retinal lactate accumulation. Retinal tissue metabolome analysis in control wild-type retina (red) and after treatment with inhibitors for MCT1 (AZD; green) or MCT1/MCT2 (ARC; blue). **a**) Principal component (PC) analysis of retinal samples investigated with ^1^H-NMR spectroscopy-based metabolomics. Dots represent individual retinal explants. **b**) Heatmap based on unsupervised hierarchical cluster analysis by Ward’s linkage, showing auto-scaled metabolite concentrations (red-high, blue-low) Squares in the heatmap represent data from individual retinal explants. **c**) Scatter plot for retinal lactate levels (arbitrary units; a.u.) and pattern hunter diagram showing top 25 correlating metabolites, correlation coefficient given as Pearson r distance. . **d**) Scatter plot for retinal ATP levels and pattern hunter. **e**) Ratios for retinal ATP/ADP and GTP/GDP levels. **f**-**h**) Scatter plots for retinal serine, taurine, and thiamine pyrophosphate (TPP) levels. n=5 retinas per condition; statistical analysis: one-way ANOVA, with Holm-Šidak post-hoc test, p-values: ****<0.0001, ***<0.001, **<0.01 *<0.05. DMG=N,N-dimethylglycine; GSSG=glutathione disulfide.

Prominent metabolic changes also concerned adenine and guanine nucleosides, among which ATP is a key indicator of cellular energy production (Figure 3d). Pattern hunter for ATP found GTP, N,N-dimethylglycine (DMG), and glutathione disulfide (GSSG) as the most correlated metabolites. Retinal DMG accumulation after AZD treatment and depletion after ARC treatment corresponds to an opposing regulation of O-phosphocholine, which could indicate an altered partitioning of choline metabolism between membrane phospholipid synthesis and mitochondrial one-carbon metabolism. The down-regulation of GSSG after MCT inhibition, while glutathione (GSH) levels remained low, is likely a consequence of decreased ATP-levels since GSH synthesis strongly depends on ATP availability^17^. The ratio between ATP and ADP may serve as further indicator of energetic status and displayed a significant down-regulation by both AZD and ARC treatment (Figure 3e). Similarly, the GTP/GDP ratio, which may reflect biosynthetic activity, is strongly decreased by MCT inhibition. Since at low ATP levels GTP can only be produced in the Krebs-cycle via GTP-forming succinyl-CoA ligase (SUCLG), low GTP-levels may be an indicator of reduced Krebs-cycle activity^11^. Remarkably, ARC treatment resulted in a significantly greater decline in both ATP/ADP and GTP/GDP ratios (Figure 3e).

Feedback inhibition of glycolysis from excess lac will lead to accumulation of glycolytic intermediates, such as 3-phosphoglycerate, which may be oxidized and then transaminated to serine. This pathway can generate additional NADH and may explain the rise in intraretinal serine levels seen after ARC treatment^18^ (Figure 3f). The decrease in threonine levels after AZD and ARC treatment may indicate its usage as metabolic fuel entering the Krebs-cycle as succinyl-CoA. The levels of the amino-sulfonic acid taurine, an antioxidant and reactive oxygen species scavenger^19,20^, were significantly increased in the ARC treated group, but not after AZD treatment. In the retina, taurine is localized mostly to photoreceptors and RPE^11^, and it is essential for photoreceptor survival^21^. Since in the ONL MCT2 targeted by ARC is expressed only in cones, the taurine changes observed may stem from altered cone metabolism. However, it is unclear whether this change relates to pathological processes or perhaps to endogenous neuroprotective mechanisms. Thiamine pyrophosphate (TPP) – an important cofactor for pyruvate-dehydrogenase and α-ketoglutarate (αKG)-dehydrogenase – behaved similar to Taurine, possibly indicating a compensatory upregulation to maintain Krebs-cycle activity.

Taken together the analysis of retinal metabolomics suggests that inhibition of lac transport severely impaired photoreceptor energy metabolism and biosynthetic processes, likely resulting in cellular dysfunction.

### MCT inhibition changes metabolite content of culture medium

Since the retinal metabolomic patterns were strongly altered by MCT inhibition, we next examined the consequent changes in the supernatant R16 culture medium. Medium samples were collected at P15 where ^1^H-NMR spectroscopy-based metabolomics identified 27 metabolites as significantly changed from control by either AZD or ARC treatment (Figure 4). Curiously, the release of lac to the medium – fresh R16 medium does not contain lac – was not affected by AZD treatment. This suggests that lac release was mediated by transporters other than MCT1, such as MCT3 expressed on RPE cells or MCT4 expressed on Müller glia cells (Supplemental Figure S1). On the other hand, ARC treatment did reduce lac release to the medium, concomitant with elevated medium glucose levels, hinting at decreased consumption and an overall reduced metabolic activity. Pattern hunter analysis for lac showed pyr, valine, and formate as the most correlated metabolites. Medium content for three metabolites was increased by AZD but decreased by ARC treatment, which could indicate transport impairment in the former and reduced metabolism in the latter case.

**Figure 4.**
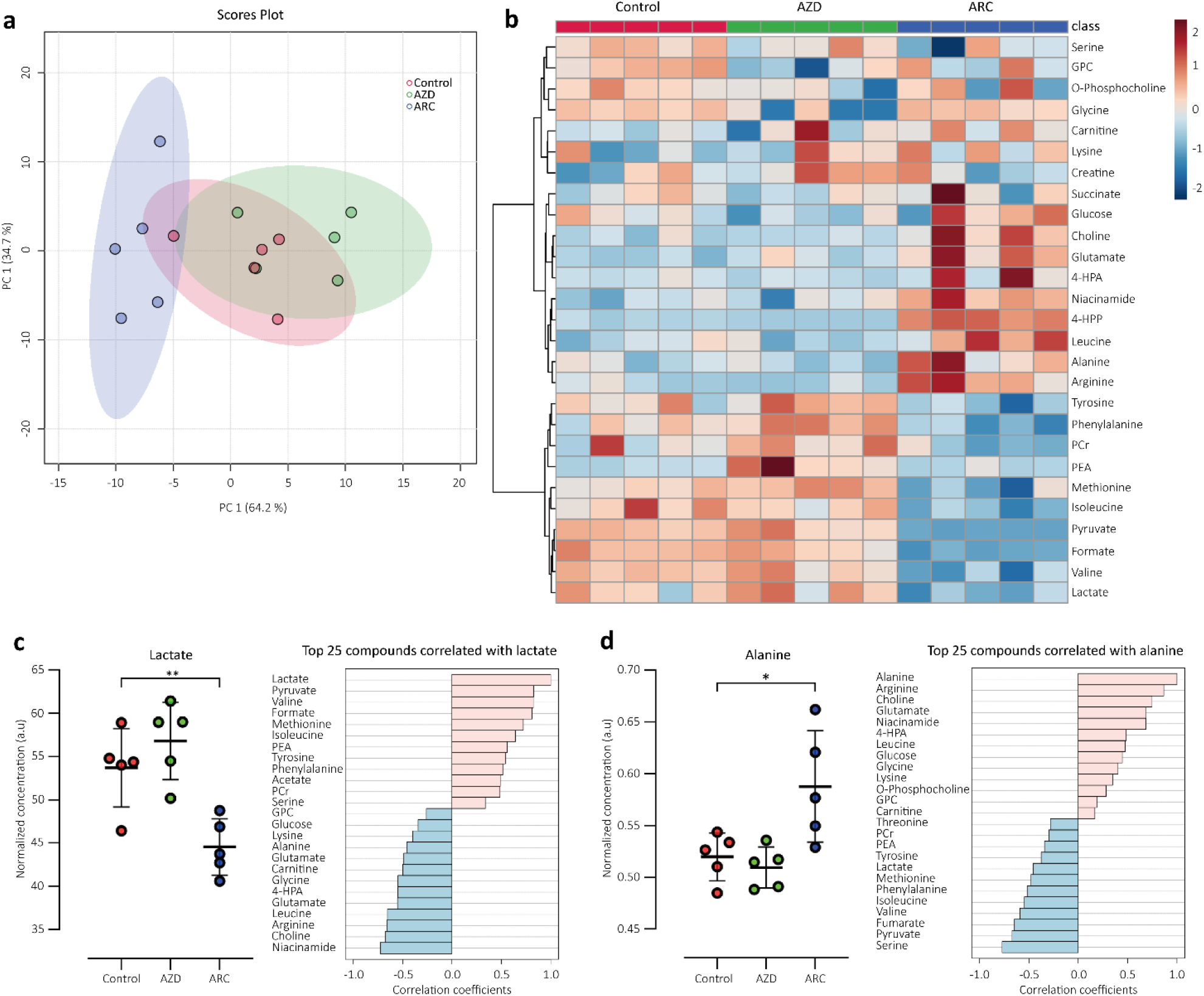
MCT1 and MCT1/MCT2 inhibition differentially alters retinal metabolite release. Analysis of metabolites released by the retina into the defined culture medium. **a**) Principal component (PC) analysis of medium samples investigated with ^1^H-NMR spectroscopy-based metabolomics. Dots represent individual medium samples. **b**) Heatmap based on unsupervised hierarchical cluster analysis by Ward’s linkage, showing auto-scaled metabolite concentrations (red-high, blue-low) in three different experimental conditions: control-red, AZD-green, ARC-dark blue. Squares represent data from individual medium samples. **c**) Scatter plot for medium lactate concentration (arbitrary units; a.u.) and pattern hunter diagram showing top 25 correlating metabolites, correlation coefficient given as Pearson r distance. **d**) Scatter plot for medium alanine concentration and pattern hunter. n=5 samples per condition; statistical analysis: one-way ANOVA, with Holm-Šidak post-hoc test; p-values: ****<0.0001, ***<0.001, **<0.01 *<0.05. 4-HPA=4-hydroxyphenylacetic acid; 4-HPP=4-hydroxyphenylpyruvate; GPC=glycerophosphocholine; PEA=O-phosphoethanolamine; PCr=creatine phosphate; TPP=thiamine pyrophosphate.

The release of alanine (Ala) to the supernatant medium after ARC treatment, could relate to compensatory Cahill-cycle activity^22^ in cones, which may use glutamate (Glu) or other amino acids as fuels in a situation where lac is no longer available^11^. However, Glu levels appear to be high in the ARC situation (see below), while a number of other amino acids are depleted (methionine, tyrosine, phenylalanine). Also, the branched-chain-amino-acids valine and isoleucine, but not leucine are strongly reduced in medium derived from ARC-treated retina. This could mean the use of valine and isoleucine as fuels, which ultimately enter the Krebs-cycle as succinyl-CoA and acetyl-CoA, respectively. Entering high levels of succinyl-CoA into the Krebs-cycle will eventually lead to a build-up of oxaloacetate (OAA), which in cones can be converted into PEP and pyr via the activities of PCK2 and PKM2^23^. The fact that leucine is apparently not used in the same way as valine and isoleucine may relate to difficulty entering its breakdown products into the Krebs-cycle.

Remarkably, the medium metabolomic patterns provide an indication for increased tyrosine aminotransferase activity after MCT inhibition: Tyrosine aminotransferase uses αKG as acceptor of tyrosinés amino group to reversibly generate 4-hydroxyphenylpyruvate (4-HPP) and Glu. In ARC medium samples tyrosine is depleted while 4-HPP and Glu levels are high. With AZD treatment this situation is reversed, indicating that here the additional generation of tyrosine and especially αKG may fuel the Krebs-cycle.

### Lactate is sufficient to maintain cone photoreceptor function

To investigate the relationship between rod and cone metabolism and their functions, we employed multi-electrode-array (MEA) recordings, to obtain localized micro-electroretinograms (µERG) from adult, P30 wild-type retina. Here, the light-evoked local field potentials reflect the electrophysiological responses of different retinal cell types, including photoreceptors and bipolar cells (BPs; 2^nd^ order neurons), while the MEA enables the simultaneous recording of retinal ganglion cell activity (RGCs; 3^rd^ order neurons)^24^. The use of scotopic (dim-light) and photopic (bright-light) stimulation allows distinguishing rod and cone photoreceptor responses, both characterized by an initial negative deflection in the µERG (aka a-wave). After µERG measurements, we performed histological workup and TUNEL labelling to ascertain that acute explantation and recording procedures did not by itself cause retinal damage. The results show that there was essentially no cell death in the ONL, while some INL cell death was likely related to regression of inner retinal vasculature (Supplemental Figure S2).

Remarkably, replacing glucose (glc) with lac as the only caloric component in the defined recording medium distinctly modulated the functions of the various neuroretinal cells: In medium containing only lac, the scotopic light response of the rod photoreceptor system (Figure 5a1, a2) displayed a 2-log-unit delay (at -2.0 light intensity) and a ≈72% reduction in the photoreceptor driven a-wave (max. amplitude: glc: - 216.2±12.2 µV; lac: -61.4±7.7 µV) when compared to recording in glc-containing medium. Strikingly, RGC activity was almost entirely abolished (Figure 5a1, a3) in lac-medium (scotopic spike count/10 ms: glc: 181.8±11.8; lac: 10.5±3.4). The MCT1-inhibitor AZD strongly reduced rod a-wave amplitudes in both glc- and lac-containing medium (Figure 5a4). Yet, treatment with the MCT1/MCT2 inhibitor ARC maintained significant rod responses in glc-medium, but not in lac-medium.

**Figure 5:**
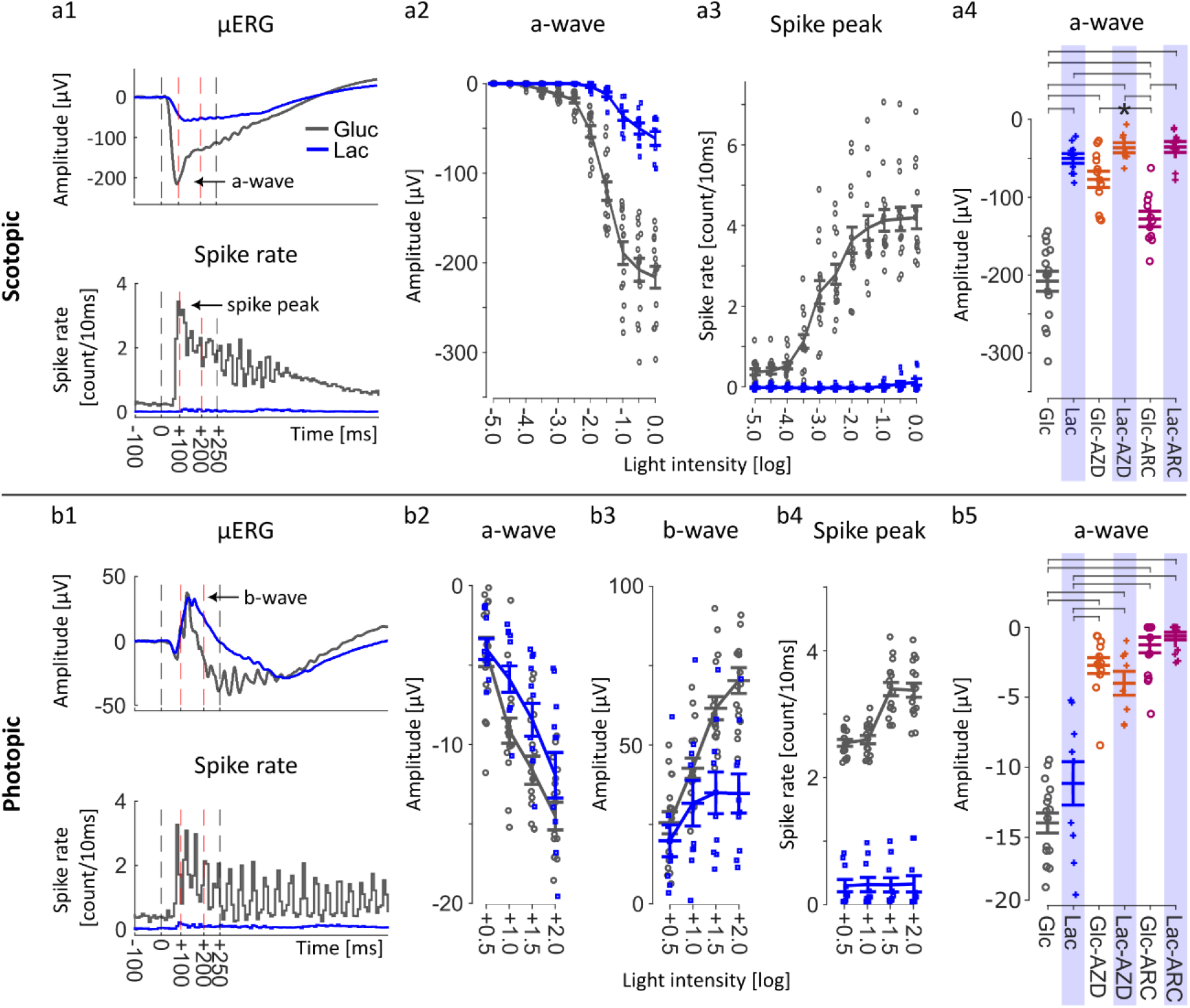
Lactate transport is critical to cone function. Retinal function was tested using micro-electroretinogram (µERG) on acutely explanted mouse retina dissected in the dark. Incubation medium contained either 20 mM glucose (glc) or 20 mM lactate (lac), with/wo inhibitors of monocarboxylate transporters (MCTs). **a**) Retinal responses under scotopic light conditions. a1) Averaged µERG recording traces obtained in medium containing glc (grey) or lac (blue) after full field 250 ms light flash with 4.20×10^13^ photons/cm²/s intensity, showing characteristic light-induced photoceptor hyperpolarization (aka a-wave). Corresponding retinal ganglion cell (RGC) responses are shown below (spike rate: spikes/10 ms bin). Spike peak indicates the 1^st^ maximum of RGC responses, correlated to maximal a-wave deflection. a2) Rod a-wave response amplitudes plotted against 11 increasing light intensities (1.33×10^9^ to 4.20×10^13^ photons/cm²/s; 0.5 log unit steps) and, a3) corresponding RGC responses. a4) Functional assessment of lactate transporters MCT1 and MCT2. Rod a-wave amplitudes in glc- or lac-containing medium with/wo MCT1 inhibitor AZD3965 (AZD; 50 µM) or MCT1/MCT2 inhibitor AR-C155858 (ARC; 20 µM). Scotopic, rod photoreceptor-driven light responses were strongly reduced by MCT inhibition, with the notable exception of ARC in glc-medium. **b**) Photopic light responses. b1) Representative photopic recordings (5 min light-adapted at background light intensity of 4.20×10^13^ photons/cm²/s; flash intensity: 4.20×10^15^ photons /cm²/s). b2) Photopic recordings with increasing light stimuli (+0.5 to +2.0; *i.e.* 1.33×10^14^ to 4.20×10^15^ photons/cm²/s). b3) Quantification of b-wave amplitudes, *i.e.* positive µERG deflections driven by bipolar cells and, b4) corresponding RGC responses. Note that photopic cone responses were preserved in lac-medium, but abolished by MCT inhibition. Number of retinal explants: Scotopic: glc: n=15, glc-AZD: n=12, glc-ARC: n=11, lac: n=10, lac-AZD: n=8, and lac-ARC: n=10. Photopic: glc: n=16, glc-AZD: n=13, glc-ARC: n=15, lac: n=10, lac-AZD: n=8, and lac-ARC: n=14; horizontal brackets indicate a significance level p<0.0001, unless indicated otherwise.

Photopic µERG responses are inherently weaker than scotopic a-waves generated by rods ^24^ since cones constitute only 3–5% of the photoreceptor population in the mouse retina ^25^. In contrast to the rod situation, when medium glc was replaced by lac, cone photoreceptor responses remained largely unaffected across all photopic light intensities (Figure 5b1, b2; a-wave: glc: -14.5±0.9 µV; lac: -11.9±1.4 µV), retaining ≈82% of the light-dependent activity. The photopic positive deflection of the µERG, the so-called b-wave, a feature driven by BP cells, was reduced by 50.6% in lac-medium (Figure 5b1, b3), when compared to recordings in glc-medium (b-wave: glc: 70.3±4.1; lac: 34.8±6.2). The photopic RGC responses – similar to what was seen at scotopic light levels – were essentially absent in lac-medium (Figure 5b1, b4; photopic spike count/10 ms: glc: 141.8±4.6; lac: 13.7±5.4). Treatment with the MCT1-inhibitor AZD strongly reduced cone a-wave amplitudes in glc-medium, but less so in lac-medium, while combined MCT1/MCT2 inhibition with ARC completely silenced cones in both glc- and lac-medium (Figure 5b5).

These findings highlighted the marked differences of various retinal cell types in their capacity to use lac as an energy source and maintain neuronal function: In the outer retina, rod function was reduced to about 1/3, indicating strong dependence on glucose. By contrast, cones could maintain their light responsiveness using lac as sole energy source. In the inner retina, BP cells retained about 50% of their functionality, while RGCs could not sustain their function at all in lac-medium. Overall, these results, notably the MCT inhibition data, lend strong support to the concept of lac-shuttling from rod to cone photoreceptors.

### Lactate maintains cone viability in inherited retinal degeneration

The very marked effect of lac on cone functionality suggested that cone metabolism was fueled to a major extent by lac, which in turn raised the question as to whether lac could also support cone viability. To address this question, we employed the *rd1* mouse model for the inherited retinal disease RP. This mouse model is characterized by a rapid loss of rod photoreceptors within the first 2-3 post-natal weeks^26^. As in human RP, *rd1* rod loss is followed by secondary cone degeneration, the reasons for which have remained enigmatic to this day.

Organotypic retinal explant cultures derived from *rd1* mice were cultivated from post-natal day 12 to 40 (28 days of culture) and exposed to either regular, serum-free and fully defined R16 medium containing 19.2 mM glc, or medium containing additional 10- or 20-mM lac. When compared to cultures derived from wild-type animals, P40 *rd1* retinal explants displayed near complete loss of rods and with that a decrease in ONL thickness to essentially one row of photoreceptors (Figure 6). Quantification of cones using immunostaining for arrestin-3 also revealed a marked cone loss in *rd1* retina. However, treatment with lac significantly increased cone numbers in a dose-dependent manner, even though lac had no apparent effect on rod degeneration.

**Figure 6.**
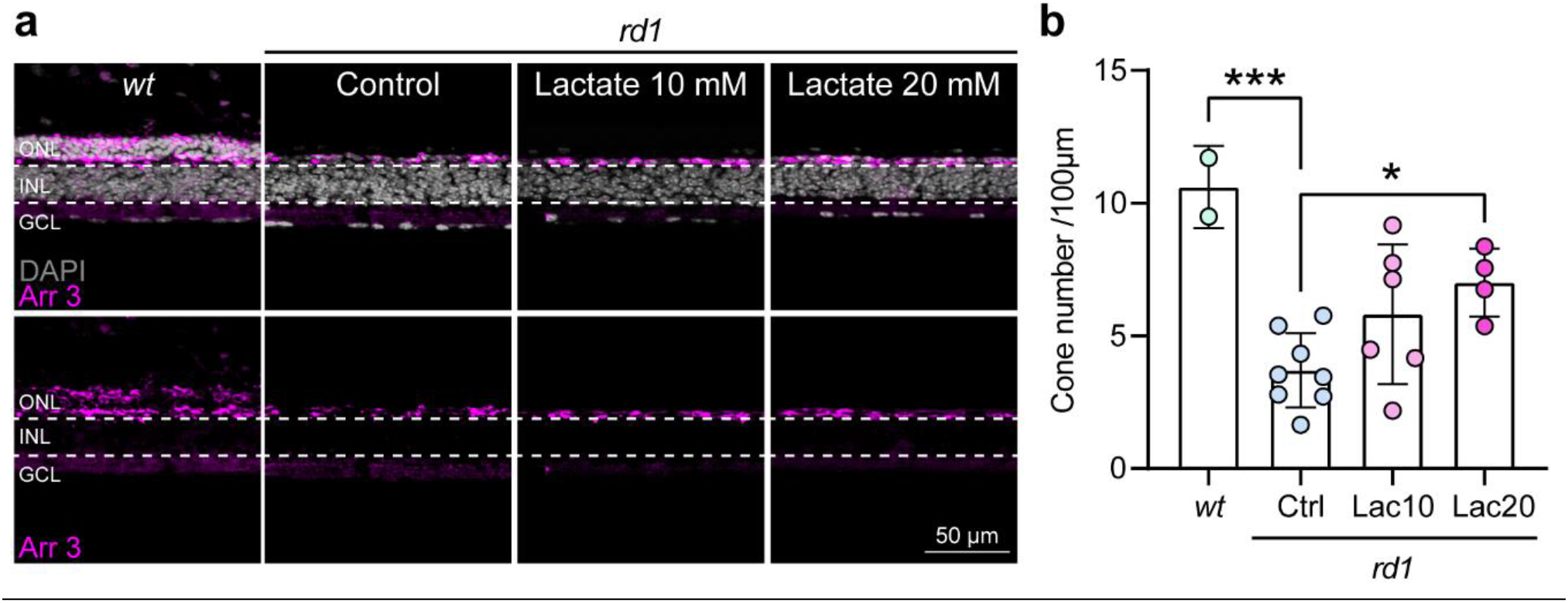
Lactate preserves cone viability in *rd1* mouse retina. **a)** Cone arrestin labelling (Arr 3, violet) in transversal sections of *rd1* retinal explants cultivated from P12 to P40, either untreated (control), or treated with lactate at 10 mM or 20 mM for 26 days. DAPI (grey) was used as a nuclear counterstain. **b)** Quantification of arrestin-positive cones in the outer nuclear layer (ONL). n=2-8 independent retinal explants. Error bars represent SD. *=p<0.05 compared to the control group using a one-way ANOVA test and Dunnett’s post-hoc test. INL=inner nuclear layer, GCL=ganglion cell layer; scale bar=50 µm.

### Mathematical simulations of rod to cone lactate shuttling in the human retina

Our data showed that cone function and viability was supported by lac released from rods. In the mouse retina rods outnumber cones approx. 20:1^25^, raising the question as to what might be the minimum number of rods required to supply one cone and whether similar lac shuttling could also apply to the human retina.

Ingram *et al*.^1^ calculated the rates at which individual mammalian rod and cone photoreceptor cells consume ATP under a range of light intensities (Figure 7a1). We used this information, together with results from the present study, to estimate individual rod and cone lac production and consumption rates. Our calculations suggest that individual rod lac production exceeds individual cone lac consumption at all light intensities (Figure 7a2). Taking the ratio of these rates, we predict that a minimum of ≈0.84 rods per cone is required to adequately sustain cone activity across all light intensities (Figure 7a3).

**Figure 7:**
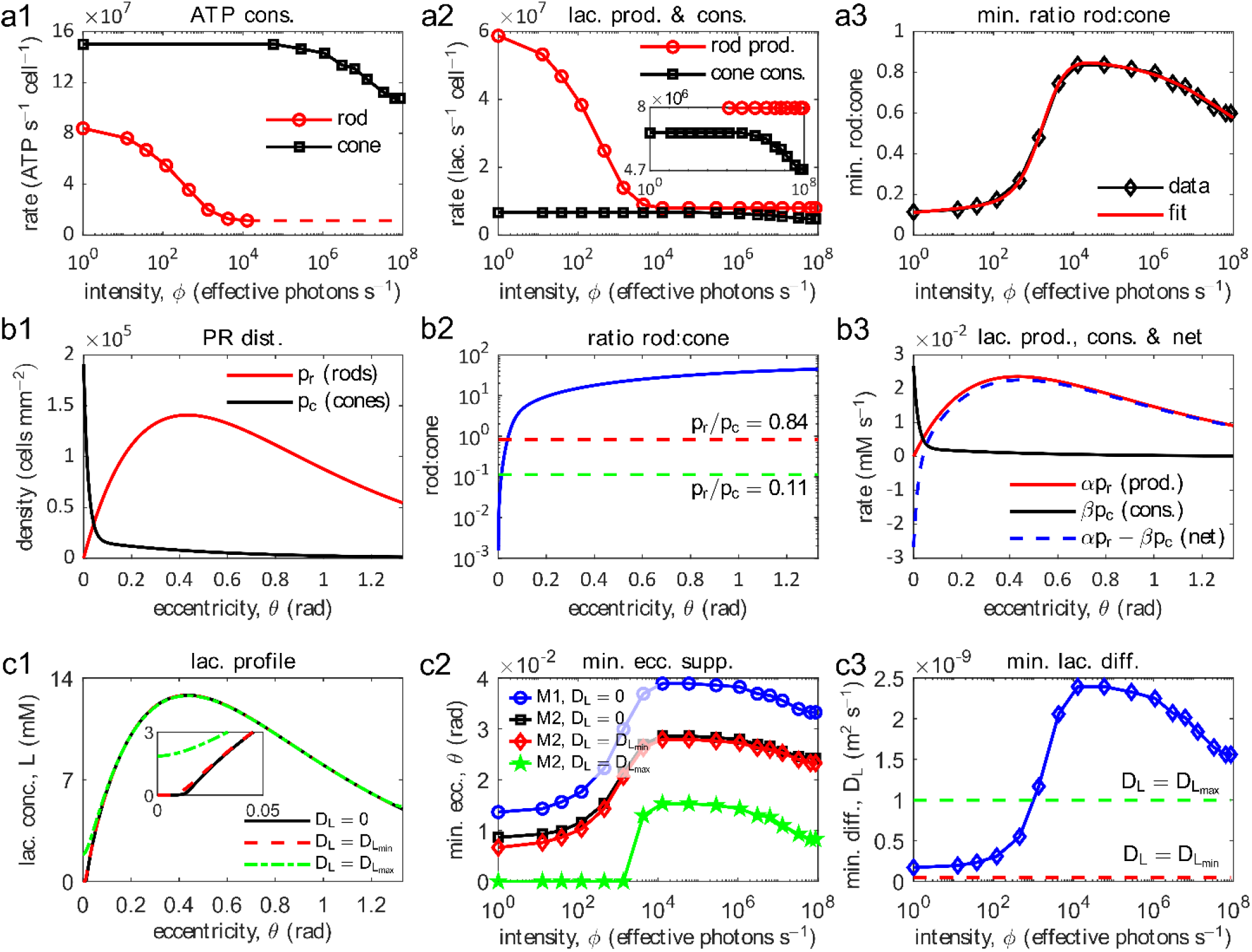
Mathematical models indicate that rods cannot supply human foveal cones with sufficient lactate. **a**) Individual cell-level calculations. a1) ATP consumption (cons.) rates of individual rod and cone cells (data from Ingram et al.^1^). a2) Lactate (lac.) production (prod.) rate of an individual rod and lac consumption rate of an individual cone (inset shows magnified region). a3) Minimum (min.) rod:cone ratio required to fully sustain cone activity. The minimum rod:cone varies with light intensity between ∼0.11 and ∼0.84. **b**) Tissue-level calculations. b1) Rod and cone photoreceptor (PR) distribution (dist.) along temporal horizontal meridian in a healthy human retina^27,28^. b2) Rod:cone (solid blue curve), and upper and lower minimum rod:cone thresholds (red and green dashed lines, respectively). b3) Gross and net rates of rod lac production and cone lac consumption (for the most challenging intensities; *φ* = 1.28 × 10^4^ to 5.90 × 10^4^ effective photons s^-1^). In the human retina, without lac diffusion, there are insufficient rods in the foveal center to sustain cone activity. **c**) Mathematical modelling of the effect of diffusion on lac supply. c1) Predicted lac concentration (conc.) profiles (Model 2, at *φ* = 1 effective photons s^-1^; inset shows magnified region). c2) The minimum eccentricity (ecc.) to which cone light-dependent activity can be fully supported (supp.; Model 1, with *η* = 0 s^-1^), or to which the rate of cone lac uptake can be maintained above 50% (Model 2, with *η* = 1.35 × 10^−2^ s^-1^). c3) The minimum lac diffusivity (diff.) required to fully sustain cone light-dependent activity across the entire retina (solid blue curve; Model 1, with *η* = 1.35 × 10^−2^ s^-1^), and lower and upper lac diffusivity bounds (red and green dashed lines respectively). Lac diffusivity would need to exceed realistic bounds in order for the centermost retina to be supplied with enough lac to fully sustain cone activity at higher illumination intensities. Parameter values: *D_Lmin_* = 5 × 10^−11^ m^2^ s^-1^ and *D_Lmax_* = 1 × 10^−9^ m^2^ s^-1^. For remaining parameter values see Table 2.

Because in the human retina cones dominate in the fovea and rods dominate in the remainder^27,28^ (Figure 7b1), the rod:cone ratio drops beneath the minimum required to support cones in a narrow central region *θ* < 3.89 × 10^−2^ rad (0.46 mm from foveal center) (Figure 7b2), with the result that the rate of cone lac consumption predicted to be required to fully sustain cone activity exceeds rod lac production in this region (Figure 7b3). However, lac diffusion from peripheral rods might be sufficient to supply this narrow central region and maintain cone function. To investigate this, we formulated two steady-state reaction-diffusion models of retinal lac distribution. Model 1 describes the theoretical scenario where cone lac consumption is maximal everywhere and Model 2 explores the scenario where cone lac consumption depends upon the local lac concentration. Diffusion increases the lac concentration in the narrow central region (Figure 7c1), increasing the proportion of the retina that can be supplied with adequate lac (Figure 7c2), and potentially meeting demand across the whole retina at lower illumination intensities (*φ* < 1.35 × 10^3^ effective photons s^-1^); however, the lac diffusivity (*D_L_* m^2^ s^-1^) would need to exceed realistic bounds to supply the whole fovea with enough lac to fully sustain cone activity at higher illumination intensities (*φ* ≥ 1.35 × 10^3^ effective photons s^-1^; Figure 7c3).

The modeling results therefore suggest that while peripheral and macular human cones can be fully supplied by rods, foveal cones cannot, suggesting that in the fovea different or additional nutrient supply mechanisms must be at play. Such a metabolic difference between foveal cones and more peripheral ones could potentially explain why central cones in both RP and in AMD initially resist the degeneration.

## Discussion

Over a hundred years ago the work of Otto Warburg raised a critical question: Why does the neuroretina resort to energy inefficient aerobic glycolysis and release large amounts of lac? Here, we show that lac produced by rod photoreceptors is not a “waste product” but a fuel that drives cone photoreceptor function and viability. The discovery of this rod-to-cone lactate shuttle may explain a number of previously enigmatic phenomena, including cone degeneration seen in retinal diseases such as RP and AMD. Moreover, lac-shuttling has important ramifications for our understanding of the CNS and may enable new forms of metabolism-centered treatments for neurodegenerative diseases.

### Photoreceptor metabolism: Use of glycolysis vs. OXPHOS

In the early 1920s Otto Warburg showed that the neuroretina, unlike any other healthy tissue, released high amounts of lac even in the presence of oxygen^29^. Since lac is generated by reduction of glycolytic pyr, Warburǵs finding implied that the retina – and by extension photoreceptors as the cell class responsible for the high retinal lac production – would resort to inefficient glycolysis to satisfy its extraordinarily high energy demand. A demand which could be satisfied far more efficiently by OXPHOS. Perhaps, even more puzzling was the fact that photoreceptors in their inner segments harbor one the highest densities of mitochondria^30^, organelles which are concerned with OXPHOS and which would scarcely be needed if photoreceptors used primarily glycolysis.

Our data on rod and cone activity in lac-only *vs*. glc-only medium, suggests that rods sustained about 30% of their light-dependent activity from OXPHOS and 70% from glycolysis, while cone activity depended to about 80% on OXPHOS and only to 20% on glycolysis. This interpretation is in line with a recent publication on rod energy metabolism^31^, yet, it rests on the assumption that no other retinal cell type employed lac for gluconeogenesis to provide glc to photoreceptors^32,33^ . While Müller glia cells have been proposed to perform gluconeogenesis and provide glc to other retinal cells in amphibian retina^34^, a direct confirmation for such a mechanism in mammalian retina seems to be lacking^2^. In our work, Müller glia gluconeogenesis is unlikely to have contributed significantly to retinal glc levels, notably since glc-dependent RGC activity was completely silenced in lac-only medium. Nevertheless, we have previously shown that lac can in part fuel the metabolism of inner retinal bipolar cells^35,36^. This corresponds to our present findings where the photopic b-wave responses were maintained in lac-only medium at approx. 50% of the response amplitudes seen in glc-medium.

While previous studies have described photoreceptor metabolism as predominantly glycolytic^9,37^, the fact that OXPHOS is essential for maintaining photoreceptor structural and functional integrity was recently confirmed by deletion of pyruvate dehydrogenase (PDH) – an enzyme involved in the formation of acetyl-CoA and the initiation of the Krebs-cycle – from either rod or cone photoreceptors^38^. In our own work, we found that in retina treated over 48 hours with the uncoupling agent Carbonyl cyanide-p-trifluoromethoxyphenylhydrazone (FCCP), rods underwent rapid degeneration associated with very strong lac production, while cones initially remained viable^11^. In view of our current data, this indicates that rods – even though most of their functional activity appears to be derived from glycolysis – require OXPHOS to maintain their viability. The resilience of cones to FCCP uncoupling may be related to the ample amounts of lac provided by rods and the activity of PCK2 and PKM2 that enables cones to produce ATP from lac even when OXPHOS is disrupted^23^ (Figure 8).

**Figure 8.**
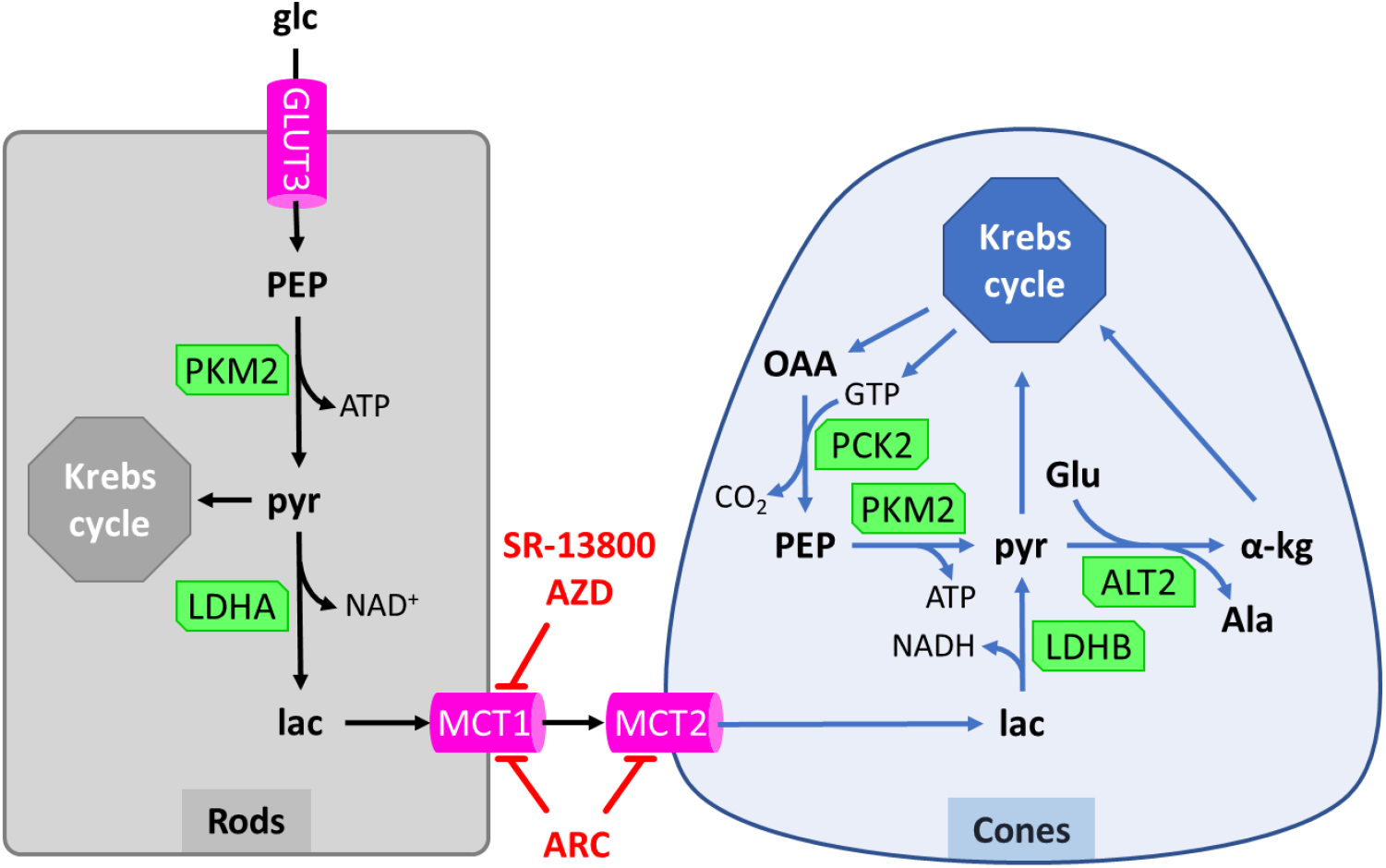
Central role of lactate shuttling in rod and cone energy metabolism. Glucose (glc) can enter photoreceptors via the high-affinity glucose transporter-3 (GLUT3). In rod photoreceptors, glc is processed to phosphoenolpyruvate (PEP), from which the enzyme pyruvate kinase M2 (PKM2) generates ATP and pyruvate (pyr). While some pyr may enter the Krebs-cycle in rods, a larger fraction is reduced by lactate dehydrogenase-A (LDHA) to create lactate (lac). Lac is then exported by rods via low-affinity monocarboxylate transporter-1 (MCT1) and imported into cones via high-affinity MCT2. The compounds SR-13800 and AZD3965 (AZD) selectively inhibit MCT1, while AR-C155858 (ARC) blocks both MCT1 and MCT2. In cone photoreceptors, lac is oxidized by LDHB to pyr, which may then enter the Krebs-cycle. Alternatively, pyr can be transaminated from glutamate (Glu) by alanine transaminase-2 (ALT2) to yield α-ketoglutarate (α-kg) and alanine (Ala). α-kg can enter the Krebs-cycle, eventually leading to an accumulation of oxaloacetate (OAA). Cones may then use pyruvate carboxykinase-2 (PCK2) and PKM2 to convert OAA back to pyr. Note that lac import and oxidation generates additional NADH, while pyr transamination can allow cones to generate ATP from amino acids.

Taken together, our data show that both rod and cone photoreceptors employ OXPHOS to satisfy their energetic needs, with cones displaying a much larger OXPHOS usage than rods, a phenomenon likely related to their significantly higher energy demand overall^1^ and corresponding to their higher inner segment mitochondrial density^30^.

### Photoreceptor lactate shuttling: Linking rod and cone metabolism

The cell-type-specific expression patterns of enzymes involved in lac production, utilization, and transport, constitute the molecular basis for lac shuttling^32,39^. Lac transport across retinal membranes is mediated by MCTs, encoded by the SLC16A gene family, which act as bidirectional proton-linked carriers of lac^40^. MCT1, MCT3, and MCT4 are primarily associated with lac export^41–43^, whereas MCT2 often functions as a neuronal lac importer^44^. Immunofluorescence localization suggested that glucose supplied by the RPE enters photoreceptors primarily through high affinity GLUT3^11^. In rods, glycolysis ends with the dephosphorylation of phosphoenolpyruvate (PEP) to pyr, a reaction catalyzed by PKM2. LDHA then reduces pyr to lac, which may subsequently be exported through MCT1 and taken up by cone photoreceptors via MCT2. Recently, genetic deletion of LDHA from rods was found to cause rod and cone degeneration^45^. In cones, LDHB can oxidize lac to pyr, generating one mole of NADH, which in turn may yield up to 3 moles of ATP via OXPHOS. Pyr may then enter the mitochondrial Krebs-cycle and OXPHOS. Alternatively, pyr and glutamate (Glu) may be transaminated by alanine aminotransferase-2 (ALT2, aka as ALAT2) to generate α-ketoglutarate (α-kg) and alanine (Ala). ALT2 was previously found to be strongly expressed in cone inner segments^11^ (*cf.* Supplemental Figure S1) and could enable cones to perform Cahill-cycle-like metabolism^22^. This will raise mitochondrial OAA levels and may be exploited for ATP-synthesis via the combined activity of PCK2 and PKM2, which will also restore pyr levels^23^ (Figure 8).

The selective inhibition of MCT1 with AZD resulted in reduced ATP production, decreased rod function, and rod degeneration, together with intraretinal lac accumulation. This suggested that MCT1 served primarily for lac export from rods and that blocking MCT1 disrupted rod energy metabolism, likely via negative feedback inhibition of glycolysis^46^. Curiously, MCT1 inhibition alone – while strongly reducing cone function – did not significantly reduce cone viability, indicative of additional lac sources, such as MCT4 expressing Müller glia cells^35^. However, combined inhibition of MCT1 and MCT2 with ARC in glc-medium caused marked degeneration of both rods and cones, suggesting that MCT2-mediated lac uptake was essential for cone survival and could not be compensated for by excess glc. ARC treatment also led to a decrease in retinal lac production and release to the surrounding medium, indicating an overall reduction in glycolytic flux and metabolic activity.

The mathematical modelling suggested that lac from less than one rod per cone is sufficient to adequately sustain cone activity across all light intensities. This, however, does not hold true for the human fovea, where even when considering lateral lac diffusion, rod lac production is unlikely to fully sustain cone function at high light intensities. This in turn implies that foveal cones must have additional/alternative fuel sources at their disposal. While evidence from human foveal Müller glia and RPE cells may currently still be scarce, we note that already macular Müller glia and RPE cells may show distinct metabolism compared to their peripheral counterparts^47,48^. Nonetheless, collectively, our data support the hypothesis of an intraretinal lac shuttle between rods and cones. Moreover, preserving rod-derived metabolic support may represent a promising therapeutic strategy to maintain peripheral and macular cone survival in retinal diseases such as RP and AMD.

### Lactate and neuroprotection of the retina

Our findings provide important insights into the pathogenesis of neurodegenerative diseases of the retina, such as retinitis pigmentosa (RP). In RP, rod photoreceptors carrying disease-causing gene mutations degenerate and die^49^. The loss of rods is followed by secondary cone degeneration, for which the underlying mechanisms are poorly understood but are thought to be driven in part by metabolic stress and nutrient deprivation^50^. Our results now allow to propose lac released from rods as a critical metabolic substrate that sustains cone function and viability.

Even further, our data may provide for a broader mechanistic framework for neuronal metabolism and neuroprotection: The work of Gerty and Carl Cori demonstrated that blood lac-levels are increased by physical exercise^51^. Together with our data on the cone neuroprotective effect of lac, this allows us to propose lac as a plausible mechanistic link between physical exercise and neuroprotection: In humans, blood lac levels show marked fluctuations, from at rest levels of ≈1 mM^52^, to 8-12 mM during exercise, to peak concentrations under strong physical activity reaching 23 mM or higher^53^. Since interphotoreceptor matrix lac-levels have been reported to be in the range of 4-13 mM^54^, and since MCTs are gradient-driven, it seems likely that physical exercise and the correspondingly high blood lac-levels will increase the availability of lac to cone photoreceptors and may thereby promote neuroprotection. Independent of its role as a metabolic fuel, lac may promote mitochondrial biogenesis and antioxidant responses in cones. In cones, in late-stage RP, a loss of the mitochondria rich inner segment (ellipsoid zone) has been observed both in animal models^55^ and in clinical studies^56^, and it is plausible to think that this loss of cone mitochondria is caused by a loss of lac supply from rods. Indeed, in rat hippocampus lac was shown to activate key regulatory factors, including PGC-1α, enhancing mitochondrial biogenesis and leading to improved oxidative capacity^57^.

Lac-mediated neuroprotection can now also explain experimental data that was previously difficult to understand: In a number of mouse models for retinal degeneration, wheel running exercises were shown to enhance cone viability^5,58–60^. Similarly, in the *rd10* mouse model for RP, environmental enrichment of the housing cages – which entices mice to increased physical activity – was shown to increase long-term cone survival and visual function^61^. In view of our data, these results may be related to increased blood lac levels resulting from physical activity of the experimental animals. Incidentally, this would be in line with the age-old idea proposed already by Plato (Timaeus 88a; *ca*. 360 B.C.E.) that physical exercise was beneficial for CNS health.

Beyond physical exercise, these studies raise the question whether it would be possible to artificially elevate lac levels for therapeutic purposes. Direct lac infusion and ocular examination have been performed in healthy human volunteers, raising blood lac levels to ≈7 mM^62^. Regrettably, in this clinical study no retinal function testing and no long-term administration was performed, leaving open the question as to whether systemic lac treatment would be beneficial in the context of retinal neurodegeneration.

As an alternative to direct infusion of lac, other therapeutic approaches may be envisioned that could result in increased availability of lac for cones. This may include the down-regulation of the histone-deacetylase Sirt6 which leads to increased expression of glycolytic regulators and glycolytic flux, leading to improved preservation of cone function in an RP mouse model^63^. A recent study showed that exogenous expression of MCT2 on both basal and apical membranes of the RPE increased cone viability in different mouse and rat RP models^64^. Given MCT2’s very low K_m_ for lac, its expression on both sides of the RPE would likely lead to uptake of lac from the blood stream and may facilitate shuttling of additional lac towards cones.

An important gap in this new understanding of lac-mediated cone protection concerns the anatomical organization of the human retina. While in most mammalian species, and in the human peripheral retina, rods outnumber cones approx. 20:1^27,65^, in the human fovea there are essentially only cones, raising the question as to how foveal cones might be supplied with lac or other energy-rich substrates. Incidentally, in RP foveal cones tend to survive the longest, leading to tunnel vision characteristic for late-stage RP^49^, and suggesting that foveal cones may have additional supply routes, or that they may use alternative metabolic fuels. For instance, lac or glutamate could be provided by foveal Müller glia cells^2,66^. Alternatively, RPE cells could provide central cones with energy substrates such as hypotaurine, which, when oxidized to taurine, could provide cones with additional NADH and ATP via the electron transport chain^11^.

Our data also warrants a note of caution on the clinical use of MCT inhibitors. Several MCT inhibitors are currently in clinical trials for the treatment of cancer^67^, including AZD3965. However, AZD3965 exhibited dose-limiting ocular toxicity, with a marked depression of ERG activity^68^, precisely in line with our µERG results.

## Conclusion

Our findings propose the existence of a retinal rod-to-cone lac shuttle, in which rods perform glycolysis and export lac via MCT1, whereas cones import lac through MCT2 and focus on OXPHOS. This intercellular lac shuttle plays a critical role in maintaining metabolic homeostasis, energy supply, and function of both photoreceptor types. Our results suggest that lac serves as an intercellular substrate that couples the metabolism of the two different neuronal cell types, presumably to allow for the most efficient substrate utilization and in ways similar to what was originally proposed for the astrocyte-neuron-lactate-shuttle hypothesis^6^. Importantly, while in the past decades the consensus in retinal degeneration research was that one had “to spare rods, to save the cones” ^78^, *i.e.* that improving rod survival would preserve cone viability and function, our work reverts this paradigm and now provides the mechanistic background and a realistic avenue for <u>direct</u> cone protection.

## Methods

### Animals

Wild type C3H mice (devoid of the *rd1* mutation) were housed under standard white cyclic illumination, had free access to food and water, and were used irrespective of gender. All personnel entrusted with animal care and handling had received appropriate training, guidance, and supervision, as specified in the German law on animal protection. The procedures were reviewed and approved by the Tübingen University review board (according to §4 German animal welfare act: AK02/19M, AK05/22M, AK02/24M), and performed in compliance with the Association for Research in Vision and Ophthalmology (ARVO) statement for the use of animals in ophthalmic and visual research. For organotypic retinal explants, mice were killed by decapitation at postnatal day (P) 9 (short-term cultures) or P12 (long-term cultures) and retinas from both eyes were used for culturing and subsequent metabolomic analysis. Adult mice, anesthetized and sacrificed in a carbon dioxide atmosphere, were used for immunostaining experiments (P30) and µERG recordings (P30-35).

### Organotypic retinal explant culture

Retinal explants obtained from either P9 wild-type mice or P12 wild-type and *rd1* mice were cultured as previously described^69^. Briefly, the retina was placed with the pigment epithelium facing down on cell culture inserts (Order No.: 83.3930.040; 0.4 µm TC-inserts; Sarstedt, Nümbrecht, Germany) with R16 complete medium (CM; order No.: 07491252A, Gibco, Paisley, UK) containing 19 mM glucose and supplements^70^, which was replaced every two days. Retinal explants were cultured in a humidified incubator at 37°C and 5% CO_2_. From P9 to P11, the cultures were kept in CM, afterwards treatments (5 µM AZD3965, 2 µM AR-C155858, or 0.1 µM SR-13800), were applied until P15. Long-term cultures of wild-type and *rd1* retina were treated with 10 or 20 mM of additional lac from P14 to P40.

### Histology, TUNEL assay, and immunofluorescence

Culturing was ended at P15 by 4% PFA fixation for 45 min at room temperature (RT). This was followed by cryoprotection in graded sucrose solutions (10%, 20%, 30%) and embedding in OCT cryometrix (Leica-Microsystems), flash-frozen on liquid nitrogen. Cryosections (12 µm) were collected on glass slides and stored at -20°C. Prior to staining, these glass slides were thawed and washed in phosphate buffered saline (PBS) for 10 min.

Terminal deoxynucleotidyl transferase dUTP nick end labeling (TUNEL) assay was performed on glass slides with retinal cryosections. After washing with PBS for 15 min, the slides were incubated with proteinase K in Tris-buffered saline (TBS) at 37 °C for 5 min. Three washes with TBS preceded treatment with ethanolic acid solution followed by three more washes with TBS, 5 min for each step. Slides were incubated in blocking buffer (PBS containing 0.3% Triton X-100, 10% normal goat serum (NGS), 1% bovine serum albumin (BSA), and 1% fish gelatin) for 1 h at room temperature in a humidity chamber. TUNEL kit (Roche Diagnostics, Mannheim, Germany) was prepared following instructions provided by supplier and slides were treated for 1 h at 37°C. Reaction was stopped by two PBS washes for 5 min each before sections were mounted with Vectashield containing DAPI (Vector Laboratories Inc., Burlingame, CA, USA) and stored in 4°C until imaging.

For immunofluorescence, blocking solution (10% NGS and 1% BSA diluted in 0.3 % PBST) was applied and slides were incubated for 1h at room temperature. Then the slides were incubated with primary antibodies diluted (Table 1) in blocking solution over night at 4°C. The following day slides were washed with PBS three times for 10 min each time. Followed by 1h incubation with corresponding secondary antibodies (Table 1), conjugated to Alexa Fluor 488 or 566 (Molecular Probes), for 1h at RT in the dark. Finally, sections were counterstained with DAPI and mounted with Vectashield (Vector Labs).

**Table 1.** Primary and secondary antibodies used in the study, providers, and dilutions.

| Antibody | Dilution | Host | Catalog # | Company |
| --- | --- | --- | --- | --- |
| GLUT1 | 1:100 | Rabbit | ab653 | Abcam |
| GLUT3 | 1:300 | Rabbit | ab41525 | Abcam |
| MCT1 | 1:100 | Rabbit | AMT-011 | Alomone labs |
| MCT2 | 1:100 | Rabbit | AMT-012 | Alomone Labs |
| MCT3 | 1:100 | Rabbit | ab60333 | abcam |
| MCT4 | 1:100 | Rabbit | ab180699 | abcam |
| LDHA | 1:100 | Rabbit | LS-C143436-100 | LSBio |
| LDHB | 1:100 | Rabbit | LS-C802610-100 | LSBio |
| PKM2 | 1:100 | Rabbit | 4053 | Cell Signaling |
| PCK1 | 1:100 | Rabbit | 7067 | Cell Signaling |
| PCK2 | 1:300 | Rabbit | NBP1-31241 | Novusbio |
| Cone arrestin (arrestin-3; Arr3) | 1:500 | Rabbit | ab15282 | Sigma |
| Peanut agglutinin (PNA), fluorescein | 1:200 |  | FL-1071 | Vector laboratories |
| Glutamine synthetase (GS) | 1:1000 | Mouse | MAB302 | Sigma-Aldrich |
| AlexaFluor568 | 1:350 | Rabbit | A11036 | Molecular Probes |
| AlexaFluor488 | 1:350 | Rabbit | A11034 | Molecular Probes |
| AlexaFluor568 | 1:350 | Mouse | A11031 | Molecular Probes |

### Microscopy and cell counting

Stained sections were analyzed using a Zeiss Z2 Apotome microscope (Carl Zeiss Microscopy, Oberkochen, Germany) and Axiovision software. The percentages of TUNEL-positive cells in INL/ONL were quantified by manually counting labeled cells in representative areas of retinal explant. The total number of cells was determined by dividing the areas evaluated through their respective average cell size in either ONL or INL. Cone photoreceptor density was quantified as number of cones per 100 µm of retinal circumference. Three retinal sections per animal (from at least three animals per condition) were evaluated. From each section, three random microscopic fields were captured, and the average value per section was used for statistical analysis.

### Metabolite extraction

After retinal explant culture, at P15, the tissue was quickly transferred into 80% methanol / 20% ethanol and snap-frozen in liquid nitrogen. A sample of the culture medium was taken from the same well plate as the retinal tissue, and snap-frozen in liquid nitrogen. Retinal tissue was placed in 400 µL of methanol (LC-MS grade), transferred to the 2 mL glass Covaris system-compatible tubes and 800 µL of methyl-tert-butyl ether (MTBE) was added, thoroughly mixed, and further subjected to metabolite extraction via ultra-sonication (Covaris E220 Evolution, Woburn, USA). After the extraction, 400 µL of ultrapure water were added for two-phase liquid separation. The aqueous phase was separated and evaporated to dryness. Similarly, 400 µL of the aqueous R16 medium sample was transferred to the 2 mL glass Covaris system-compatible tube, 800 µL of MTBE was added and subjected to the ultra-sonication extraction protocol. Finally, 400 µL of methanol were added and mixed, centrifuged, and after two-phase separation the aqueous layer was separated and evaporated, to obtain a dry metabolite pellet.

### 1H-NMR spectroscopy measurements and data analysis

Dried metabolite pellets were resuspended in a deuterated phosphate buffer (pH corrected for 7.4) with 1 mM of 3-(trimethylsilyl) propionic-2,2,3,3-d_4_ acid sodium salt (TSP) as internal standard. R16 medium samples did not use TSP standard. NMR spectra were recorded at 298 K on a 14.1 Tesla ultra-shielded NMR spectrometer at 600 MHz proton frequency (Avance III HD, Bruker BioSpin, Ettlingen, Germany) equipped with a triple resonance 1.7 mm room temperature micro probe. Short zero-go (zg), 1D nuclear Overhauser effect spectroscopy (NOESY) and Carr-Purcell-Meiboom-Gill (CPMG; 4096 scans for retinal tissue samples, 128 for medium samples) pulse programs were used for spectra acquisition. Spectra were processed with TopSpin 3.6.1 software (Bruker BioSpin).

Retinal tissue metabolite assignment and quantification was done on the pre-processed CPMG spectra and performed with ChenomX NMR Suite 8.5 (Chenomx Inc., Edmonton, Canada). For absolute quantification, metabolite matches were calibrated to the internal 1 mM TSP standard. After manual assignment and quantification of each sample, a raw metabolite concentration table was exported and used as upload file for statistical analysis with MetaboAnalyst 6.0 platform (https://www.metaboanalyst.ca). Raw metabolite concentrations were normalized by the probabilistic quotient normalization (PQN) method on the control group to account for dilution effects. Heat maps display auto-scaled metabolite concentrations (red-high, blue-low). Unsupervised hierarchical cluster analysis was performed by Euclidean distance measure and Ward distance measure.

Pattern hunter correlation analysis was performed on respective datasets in each specific comparison of selected metabolites based on the most statistically significant changes and importance using Pearson r distance measure for correlation coefficient calculation. GraphPad Prism 10.1.1 software (GraphPad Software, San Diego, CA, USA) was used for statistical analysis and data visualization. For multiple group comparisons an ordinary one-way ANOVA with Holm-Šidak multiple comparisons post-hoc test was applied. Fold change (FC) threshold was set to > 1.2, and raw p value < 0.05, *p*-values represent: **** < 0.0001, *** < 0.001, ** < 0.01 * < 0.05. Data throughout is represented as individual data points with mean ± SD. Each mean represents at least 5 individual biological replicates (*i.e.* retinal explant cultures from different animals).

### µERG in retinal explants

Retinal explant µERG recordings were performed as established and described previously^24^. Before the experiments, the mice were kept for at least 12 h in an air-ventilated, light-tight box for dark adaptation. Acute retinal explants were prepared out under dim red-light. The eyes were enucleated immediately after euthanasia. The incubation, dissection, and recording medium was carbonated (95% CO_2_/5% O_2_) artificial cerebrospinal fluid (ACSF). ACSF solution contained either glucose [20 mM] or lactate [20 mM] with lactate transporter inhibitors (MCT1 inhibitor: AZD3965 (AZD) 50 µM; MCT1/MCT2 inhibitor AR-C155858 (ARC) 20 µM). To assess the light-dependent retinal responses (µERG and RGC spikes), electrophysiological recordings were performed using a multi-electrode array (MEA, Multi Channel Systems (MCS), Reutlingen, Germany) system.

To discriminate rod and cone photoreceptor responses, light-stimulation and µERG recording was carried out under scotopic (rod photoreceptor responses; *in vivo* mouse dark-adaptation > 12 h) and photopic conditions (cone photoreceptor responses; *ex vivo* light-adapted). Subsequent to scotopic stimulation, the retinal explants were light-adapted on the MEA for 5 min at an intensity of 4.20 × 10^13^ photons/cm²/s. The light stimulus parameters were set as a 250 ms light flash (3 repetitions per light intensity, 20 s intervals). For scotopic conditions, there were 11 stimuli in 0.5 log steps ranging from −5.0 to 0.0 (1.33 × 10^9^ to 4.20 × 10^13^ photons/cm²/s), where 0.0 intensity corresponded to the level of photopic adaptation and background illumination for photopic recordings. For photopic conditions (background light: 4.20 × 1013 photons/cm²/s), there were four stimuli in 0.5 log steps ranging from +0.5 to +2.0 (1.33 × 10^14^ to 4.20 × 10^15^ photons /cm²/s). The light intensity was set using neutral density filters.

### Data analysis, statistics, figure preparation

Data were analyzed using Microsoft Excel (Microsoft Corporation, Redmond, WA, USA) and GraphPad Prism 11.0.2 software (GraphPad Software, San Diego, CA, USA). Values are given as mean ± SD. Statistical tests included 2-tailed Student’s t-test for two groups or one-way ANOVA for multiple groups where appropriate. Significance levels as indicated by asterisks were: * p < 0.05; **p < 0.01, *** p < 0.001, ****p < 0.0001. µERG data evaluation was carried out using custom-written MATLAB scripts (R2024a, MathWorks, Natick, USA). Statistical significance was estimated using one-way ANOVA followed by Dunnett’s multiple comparisons test. Experimental data are given as mean ± SD. Figures were prepared using Inkscape (inkscape.org, v1.2, accessed on 16 June 2024).

### Mathematical modelling of retinal lactate production, consumption, and diffusion

#### Individual cell-level

Ingram *et al*.^1^ calculated the rates at which individual mammalian rod and cone photoreceptor cells consume ATP under a range of light intensities (Figure 7a1; data extracted from Ingram et al.^1^ Figure 2C, using plotdigitizer.com). We denote these rates as *ξ_r_*(*φ*) ATP s^-1^ cell^-1^ (rods) and *ξ_c_*(*φ*) ATP s^-1^ cell^-1^ (cones), noting that they are functions of light intensity, *φ*, where 1 ≤ *φ* ≤ 10^8^effective photons s^-1^. The present study suggests that rods sustain about 70% of their light-dependent activity from glycolysis, while cones sustain about 80% of their light-dependent activity from OXPHOS. We notate these (dimensionless) proportions as *ρ_rgly_* = 0.7 and *ρ_coxp_*_ℎ*os*_ = 0.8, respectively. For every lactate molecule produced by rods following glycolysis, one ATP molecule is produced during glycolysis, while each lactate molecule can be used to produce 18 ATP molecules by cones through OXPHOS. We denote these quantities as *A_gly_* = 1 ATP lactate^-1^ and *A_oxp_*_ℎ*os*_ = 18 ATP lactate^-1^, respectively. Hence, we can calculate the rate of lactate production by rods as *ρ_rgly_ξ_r_*(*φ*)⁄*A_gly_* lactate s^-1^ cell^-1^ and the rate of lactate consumption by cones as *ρ_coxp_*_ℎ*os*_ *ξ_c_*(*φ*)⁄*A_oxp_*_ℎ*os*_ lactate s^-1^ cell^-1^ (Figure 7a2). Therefore, in order for cones to be supplied with sufficient lactate to fully sustain their activity, the following inequality must be satisfied: 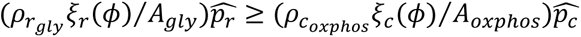 lactate s^-1^, where 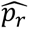 cells and 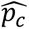 cells denote local rod and cone numbers respectively. Thus, we can calculate the (dimensionless) minimum number of rods per cone (or minimum rod:cone) required to fully sustain cone activity as follows:

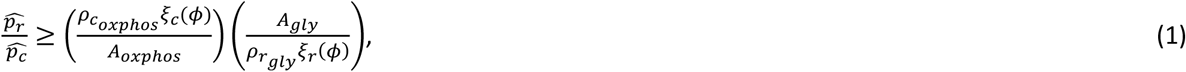

as a function of the light intensity, *φ* (Figure 7a3).

To allow for interpolation and for future convenience, we fit the following rational algebraic function to the right-hand side of Equation (1) using the Levenberg-Marquardt algorithm within MATLAB R2024b’s curve fitter toolbox:

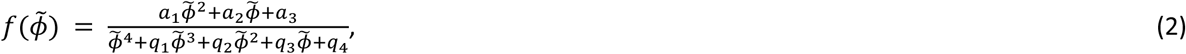

where Φ̃ = log_10_(*φ*), and the (dimensionless) parameter values are as follows: *a*_1_ = 8.54, *a*_2_ = −46.1, *a*_3_ = 73.2, *q*_1_ = −19.2, *q*_2_ = 150, *q*_3_ = −516 and *q*_4_ = 661 (to three significant figures, Figure 7a3).

### Tissue-level

To explore the extent to which rod lactate production can meet cone lactate demand in the human retina, we must account for in the human retina’s heterogenous rod and cone distributions. To do this, we utilize the following rod and cone density functions developed in Roberts *et al*. ^28^

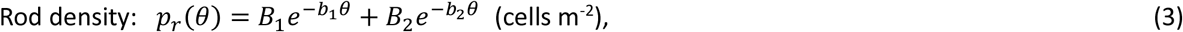

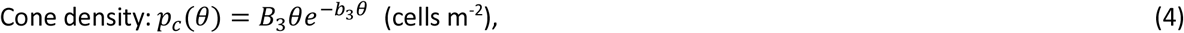

where *θ* is the retinal eccentricity (or polar angle) which runs from *θ* = 0 rad at the foveal centre to *θ* = Θ = 1.33 rad at the ora serrata. The density functions were fitted to the healthy human rod and cone distributions measured along the temporal horizontal meridian by Curcio et al.^27^ (from which we also derive our value for Θ; see Table 2 for parameter values and Figure 7b1).

**Table 2.** Parameters used in Equations (3) - (7).

| Parameter | Description | Value | Source |
| --- | --- | --- | --- |
| $R$ | Retinal radial position | $1.2 \times 10^{-2}$ m | 75 |
| $\Theta$ | Eccentricity of the ora serrata | 1.33 rad | 27 |
| $D_L$ | Lactate diffusivity | $5 \times 10^{-11}$ to $1 \times 10^{-9}$ $\text{m}^2 \text{s}^{-1}$ | 76,77 |
| $\alpha$ | Rate constant of lactate production by rods | $1.68 \times 10^{-13}$ to $1.24 \times 10^{-12}$ $\text{mM s}^{-1} \text{m}^2 \text{cell}^{-1}$ | 1,71 |
| $\beta$ | Rate constant of (maximal) lactate consumption by cones | $1 \times 10^{-13}$ to $1.4 \times 10^{-13}$ $\text{mM s}^{-1} \text{m}^2 \text{cell}^{-1}$ | 1,71 |
| $\gamma$ | Lactate concentration at which the cone lactate consumption rate is half maximal | $1 \times 10^{-2}$ mM | Estimate* |
| $\eta$ | Rate constant of cone-independent lactate consumption/loss | $1.35 \times 10^{-2}$ $\text{s}^{-1}$ | Estimated** based on 54 |
| $B_1$ | Cone profile parameter | $1.73 \times 10^{11}$ $\text{cells m}^{-2}$ | Fitted using data from 27 |
| $B_2$ | Cone profile parameter | $1.76 \times 10^{10}$ $\text{cells m}^{-2}$ | Fitted using data from 27 |
| $B_3$ | Rod profile parameter | $8.84 \times 10^{11}$ $\text{cells m}^{-2} \text{rad}^{-1}$ | Fitted using data from 27 |
| $b_1$ | Cone profile parameter | $54.1 \text{ rad}^{-1}$ | Fitted using data from 27 |
| $b_2$ | Cone profile parameter | $2.01 \text{ rad}^{-1}$ | Fitted using data from 27 |
| $b_3$ | Rod profile parameter | $2.31 \text{ rad}^{-1}$ | Fitted using data from 27 |

\*Chosen to ensure a near-maximal cone lactate consumption rate across all except the most central retinal regions. \*\*Chosen to achieve realistic retinal lactate concentrations.

The variation in the rod:cone with eccentricity can then be calculated as *p_r_*(*θ*)⁄*p_c_*(*θ*) = (*B*_1_*e*^−*b*1*θ*^ + *B*_2_*e*^−*b*2*θ*^)⁄(*B*_3_*θe*^−*b*3*θ*^) (Figure 7b2).

Starting with the rate of lactate production by a single rod, *ρ_rgly_ξ_r_*(*φ*)⁄*A_gly_* lactate s^-1^ cell^-1^, we convert this to the rate constant of lactate production by rods, *α* mM s^-1^ m^2^ cell^-1^, by dividing by the Avogadro constant (*N_A_* = 6.02 × 10^23^ molecules mol^-1^) to give units of mol s^-1^ cell^-1^, and multiplying by the surface area to volume ratio of the region being modelled (1.27 × 10^4^ m^-1^, assuming that the lactate being produced and consumed by rods and cones is largely confined to the inner segment and outer nuclear layers, which we take to lie between 46.5 *μ*m and 126 *μ*m from the choroid^71–73^ to give units of mol m^-1^ s^-1^ cell^-1^ (which can be rewritten as mM s^-1^ m^2^ cell^-1^). When multiplied by the rod density function, this gives the local rate of rod lactate production *αp_r_*(*θ*) mM s^-1^. In the same way, we can convert the rate of lactate consumption by a single cone, *ρ_coxp_*_ℎ*os*_ *ξ_c_*(*φ*)⁄*A_oxp_*_ℎ*os*_ lactate s^-1^ cell^-1^, to the rate constant of (maximal) lactate consumption by cones, *β* mM s^-1^ m^2^ cell^-1^, which, when multiplied by the cone density function, gives the (maximal) rate of cone lactate consumption *βp_c_*(*θ*) mM s^-1^. Thus, we can predict how lactate production and consumption varies with eccentricity, together with the net rate of production/consumption as *αp_r_*(*θ*) − *βp_c_*(*θ*) (Figure 7b3). We note that, while we omit the *φ* for notational simplicity, both *α* and *β* depend upon *φ* (i.e., *α* = *α*(*φ*) and *β* = *β*(*φ*)), such that the absolute (*α*(*φ*)*p_r_*(*θ*) and *β*(*φ*)*p_c_*(*θ*)) and net (*α*(*φ*)*p_r_*(*θ*) − *β*(*φ*)*p_c_*(*θ*)) rates of lactate production and consumption depend upon both retinal eccentricity and illumination intensity.

### Diffusive Effects

Finally, we examine the effects of diffusion in meeting cone lactate demand. To this end we formulate two steady-state reaction-diffusion models of retinal lactate distribution, taking a similar approach to that in Roberts et al.^28^ and Roberts^74^ (Figure 7c1). The models are posed in a spherical polar coordinate system, with origin at the posterior segment center and *z*-axis directed outward through the foveal centre. For simplicity, we neglect variation in the azimuthal direction (which accounts for rotation about the *z*-axis) and depth-average through the retina, assuming a fixed radius of *R* m (noting that the retinal width is approximately two orders of magnitude smaller than its radius of curvature). This reduces the models to a single spatial dimension (1D), this being the eccentricity, or polar angle, *θ*, which runs from *θ* = 0 rad at the foveal centre to *θ* = Θ = 1.33 rad at the ora serrata. Both models describe the steady-state lactate concentration, *L*(*θ*) mM, where Model 1 describes the (theoretical) scenario where cone lactate consumption is maximal everywhere and Model 2 explores the scenario where cone lactate consumption depends upon the local lactate concentration.

The Model 1 governing equation takes the following form:

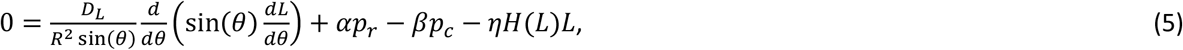

where the first term on the right-hand side accounts for lactate diffusion, with diffusivity *D_L_* m^2^ s^-1^, the second and third terms describe rod lactate production and cone lactate consumption (as described above), with *p_r_* and *p_c_* given by Equations (3) and (4), and the final term accounts for cone-independent lactate consumption/loss (e.g., consumption by RPE or Müller glia cells, or loss to the choroid), with rate constant *η*s^-1^. Since *βp_c_* exceeds *αp_r_* close to the foveal centre, this model permits negative lactate concentrations when diffusion is unable to meet demand (that is, when the magnitude of *D_L_* is insufficient). While negative lactate concentrations are unphysical, this model is useful as it allows us to determine the minimum eccentricity to which cone light-dependent activity can be fully supported (Figure 7c2) and to calculate the minimum lactate diffusivity, *D_L_*, required to fully sustain cone light-dependent activity across the entire retina (i.e., to maintain *L*(*θ*) ≥ 0, for 0 ≤ *θ* ≤ Θ; see Figure 7c3). The final term contains a Heaviside step function, *H*(*L*), which is defined such that

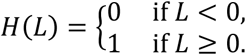

This ensures that the consumption term does not become a production term in the case that *L* < 0. The Model 2 governing equation takes the following form:

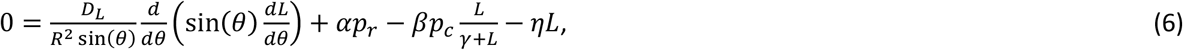

where the first and second terms on the right-hand side are the same as in Model 1 (Equation 5). The third term, accounting for cone lactate consumption, is modified to depend upon the local lactate concentration through multiplication by a Michaelis-Menten function, *L*⁄(*γ* + *L*), where *γ* mM is the lactate concentration at which lactate consumption is half maximal. Since the inclusion of the Michaelis-Menten function prevents the lactate concentration from becoming negative, the Heaviside step function is not required in the final term. Model 2 allows us to explore the minimum eccentricity to which the rate of cone lactate uptake can be maintained above a given level (e.g., *L*⁄(*γ* + *L*) ≥ 0.5; Figure 7c2).

We close Models 1 and 2 by applying zero-flux boundary conditions:

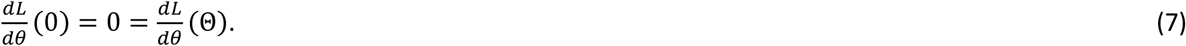

These conditions are justified by symmetry at the foveal center (*θ* = 0 rad) and by the fact that the retina terminates at the ora serrata (*θ* = Θ rad).

Models 1 and 2 were solved numerically using MATLAB R2024b’s fsolve routine, employing the trust-region dogleg algorithm, using either 1001 or 4001 mesh points. The parameter values associated with Equations (3)—(7) are given in Table 2.

## Acknowledgements

The authors would like to thank Norman Rieger for excellent technical assistance and Thomas Euler and Mathias Seeliger for expert advice (all from Institute for Ophthalmic Research, Eberhard-Karls-Universität Tübingen, Germany. Thanks for expert advice also go to James B. Hurley (University of Washington, Seattle, WA, USA) and Daniel T. Hass (Emory University Atlanta, GA, USA). We would also like to acknowledge Gordon Fain and Sampath Alapakkam for providing the original dataset on ATP consumption in rods and cones (from Ingram *et al*., Proc Natl Acad Sci U S A. 117, 19599-19603, 2020).

## Additional datasets

**Supplemental Figure S1.**
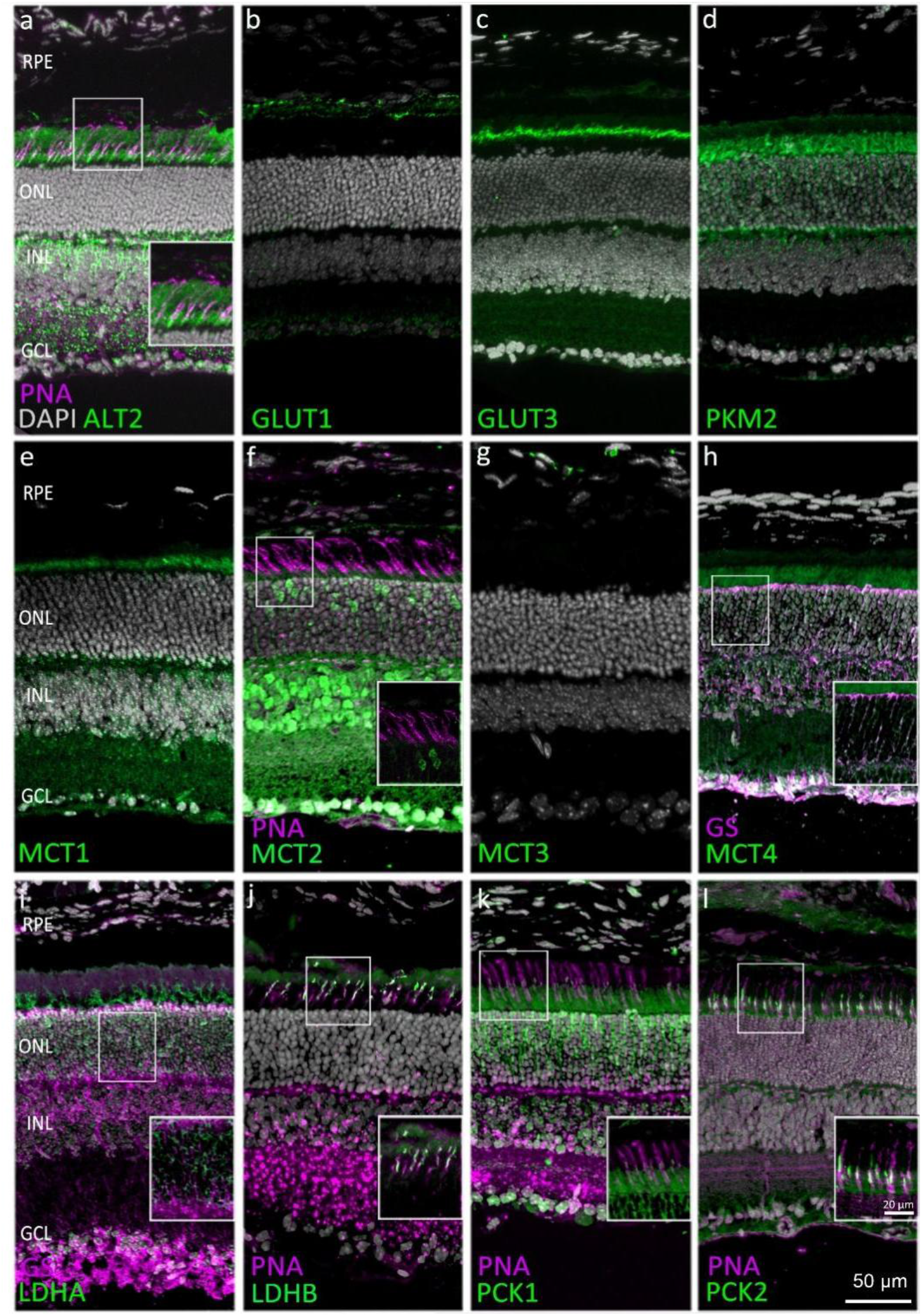
Expression of key enzymes and transporters involved in lactate metabolism. Immunodetection of key metabolic (green) enzymes molecules at post-natal day (P) 30 in wild-type retina. DAPI (grey) was used as nuclear counterstain. Co-localization with Müller cell marker glutamine synthetase (GS; magenta) and cone photoreceptor marker peanut agglutinin (PNA; magenta) was used to confirm cell-type specific expression. **a**) Alanine aminotransferase 2 (ALT2) is located in the photoreceptor segments. **b, c)** Glucose transporters -1 and -3 (GLUT1; GLUT3) are expressed in retinal pigment epithelium and photoreceptor segments, respectively. **d)** Pyruvate kinase M2 (PKM2) localizes to photoreceptor inner segments. **e-h)** Monocarboxylate transporters-1 to -4 (MCT1-4) mediate lactate transport. **i, j)** Lactate dehydrogenases, catalyze the interconversion between lactate and pyruvate. LDHA promotes lactate production, LDHB facilitates lactate oxidation. **k, l)** Phosphoenolpyruvate carboxykinase -1 and -2 catalyze the decarboxylation and phosphorylation of oxaloacetate to form phosphoenolpyruvate. ONL = outer nuclear layer, INL = inner nuclear layer, GCL = ganglion cell layer; scale bar = 50 µm.

**Supplemental Figure S2.**
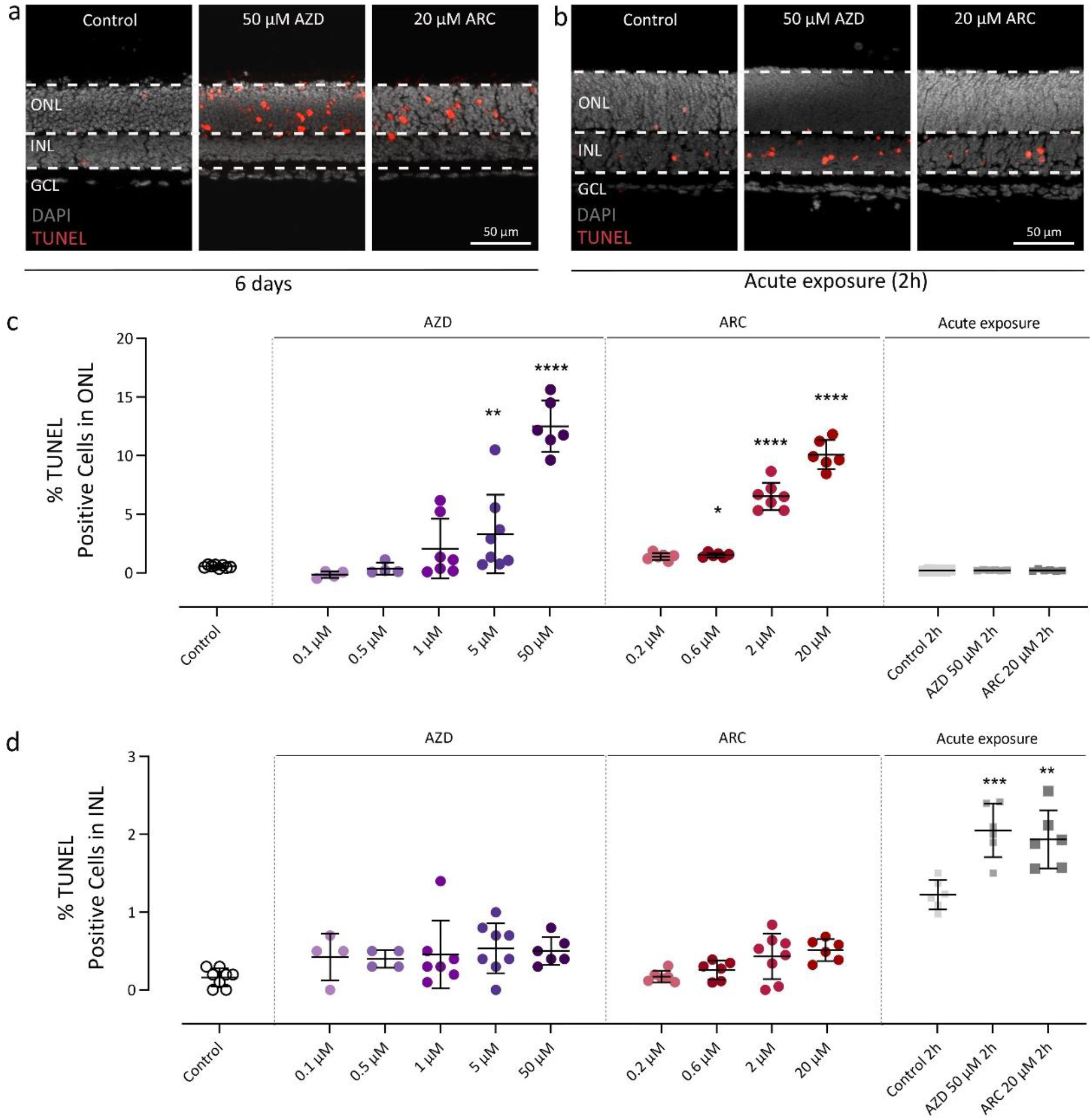
Retinal dose response properties of MCT inhibitors. Organotypic retinal explants were derived from wild-type mice at post-natal day (P) 9 and cultured for 6 days *in vitro*. From P11 onwards retinal cultures were treated with rising concentrations of the MCT inhibitors AZD3965 or AR-C155858. **a**) The TUNEL assay (red) was used to label dying cells, which were then quantified in both outer nuclear layer (ONL) and inner nuclear layer (INL). DAPI (grey) was used as nuclear counterstain. **b**) In separate experiments, the effect of the highest inhibitor concentration was assessed on acutely explanted P15 retina after 2-hour incubation. **c**) Quantification of TUNEL positive, dying cells in the ONL after long-term (4 days) and acute (2 hours) exposure. **d**) Quantification of cell death in the INL after long-term and acute exposure. n=4-8 independent retinal explants; error bars represent SD; significance levels: * = p < 0.05; ** = p < 0.01; *** = p < 0.001; **** = p < 0.0001. GCL = ganglion cell layer; scale bar = 50 µm.

**Supplemental Figure S3.**
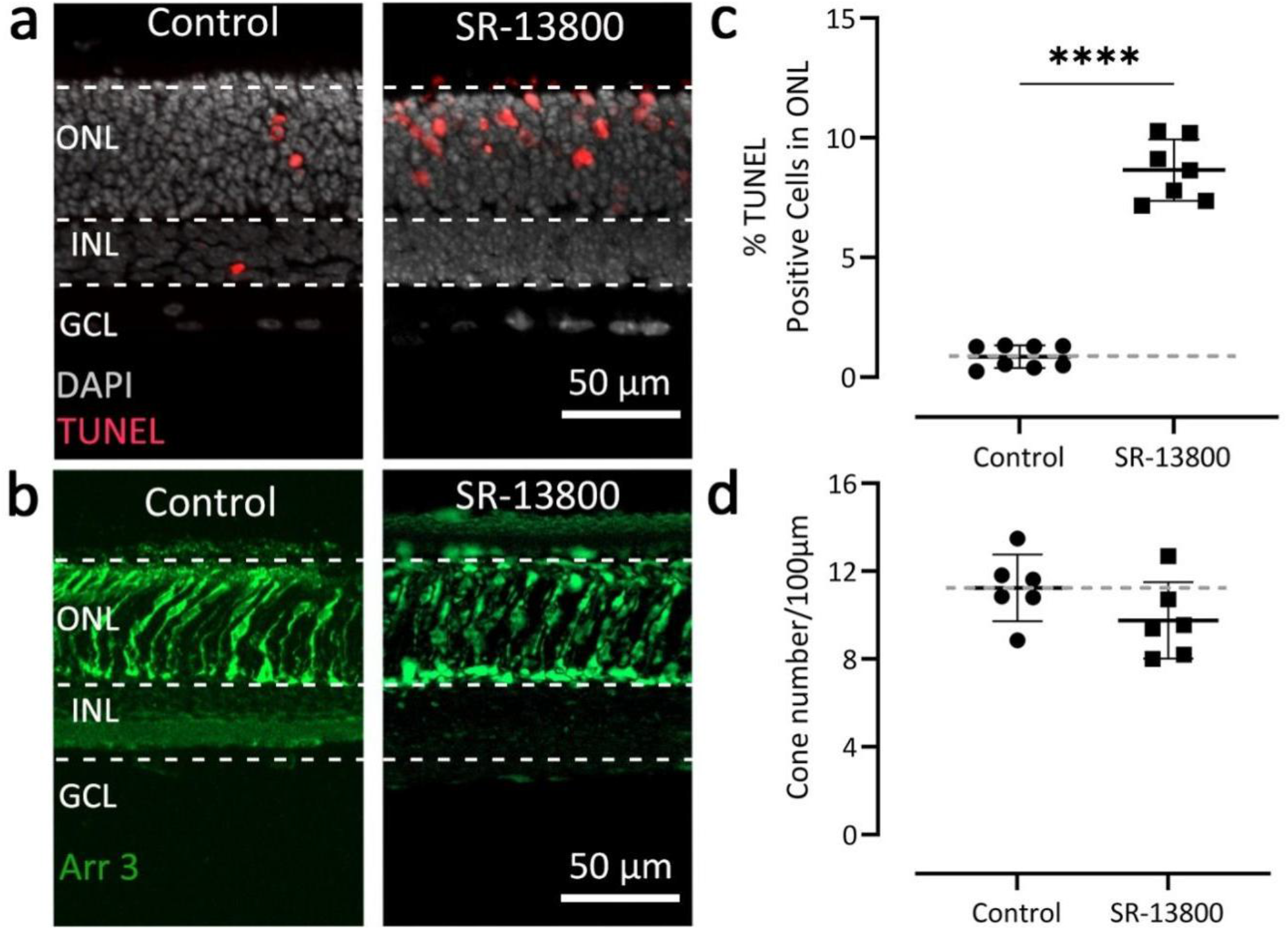
The MCT1-specific inhibitor SR-13800 selectively destroys rod photoreceptors. Organotypic retinal explants were derived from wild-type mice at post-natal day (P) 9 and cultured for 6 days *in vitro*. From P11 onwards retinal cultures were treated with 0.1 µM of the MCT1 inhibitor SR-13800. **a**) The TUNEL assay labelled dying cells in the outer nuclear layer (ONL). **b**) Immunostaining for arrestin-3 was used to label cone photoreceptors. **c**) Quantification of TUNEL positive cells shows a strong increase of cell death in the ONL induced by SR-13800. **d**) Treatment with SR-13800 did not cause significant loss of cone photoreceptors. Dashed grey lines indicate control levels; n=6-8 independent retinal explants; error bars represent SD; significance levels: **** = p < 0.0001. RPE = retinal pigment epithelium, GCL = ganglion cell layer; scale bar = 50 µm.

